# An uncanonical CDK6 activity inhibits cilia function by suppressing axoneme polyglutamylation

**DOI:** 10.1101/2023.06.22.546046

**Authors:** Kai He, Xiaobo Sun, Chuan Chen, San Luc, Jielu Hao, Yingyi Zhang, Yan Huang, Haitao Wang, Kun Ling, Jinghua Hu

**Affiliations:** Department of Biochemistry and Molecular Biology, Mayo Clinic, Rochester, Minnesota, USA; Division of Nephrology and Hypertension, Mayo Clinic, Rochester, Minnesota, USA; Mayo Clinic Robert M. and Billie Kelley Pirnie Translational Polycystic Kidney Disease Center, Mayo Clinic, Rochester, Minnesota, USA; Department of Physiology and Biomedical Engineering, Mayo Clinic, Rochester, Minnesota, USA.; Mayo Clinic Robert and Arlene Kogod Center on Aging, Mayo Clinic, Rochester, Minnesota, USA.; Division of Geriatric Medicine & Gerontology, Department of Medicine, Mayo Clinic, Rochester, Minnesota, USA

**Keywords:** CDK6/Ciliopathy/FIP5/Polyglutamylation/Primary cilia

## Abstract

Tubulin polyglutamylation is a post-translational modification that occurs primarily along the axoneme of cilia. Defective axoneme polyglutamylation impairs cilia function and has been correlated with ciliopathies, including Joubert Syndrome (JBTS). However, the precise mechanisms regulating proper axoneme polyglutamylation remain vague. Here, we show that Cyclin-Dependent Kinase 6 (CDK6), but not its paralog CDK4, localizes to cilia base and suppresses axoneme polyglutamylation by phosphorylating RAB11-interacting protein FIP5 at site S641, a critical regulator of cilia import of glutamylases. S641 phosphorylation disrupts the ciliary recruitment of FIP5 by impairing its association with RAB11, thereby reducing the ciliary import of glutamylases. Interestingly, significant upregulation of CDK6 and defective axoneme polyglutamylation were observed in Autosomal dominant polycystic kidney disease (ADPKD) cells. Encouragingly, the FDA-approved CDK4/6 inhibitor Abemaciclib can effectively restore cilia function in JBTS and ADPKD cells with defective glutamylation and suppresses renal cystogenesis in an *ex vivo* ADPKD model. In summary, our study elucidates regulatory mechanisms governing axoneme polyglutamylation and suggests developing CDK6-specific inhibitors could be a promising therapeutic strategy to enhance cilia function in ciliopathy patients.

## Introduction

Microtubule-based primary cilium senses and transduces environmental cues to diverse cell signalings essential for development and tissue homeostasis (Anvarian, Mykytyn et al., 2019, Goetz & Anderson, 2010). Dysfunctional cilia lead to dozens of rare genetic disorders, (e.g. ADPKD, Joubert syndrome), which collectively termed ciliopathies (Reiter & Leroux, 2017). In general, the severity of ciliopathies depends on the nature of the genetic mutation to what extent it affects cilia function. Restoration of defective cilia function in ciliopathies is a promising but challenging therapeutic strategy due to lack of mechanistic understanding of pathogenesis for most ciliopathies and, also, druggable targets with safe profiles.

Tubulin posttranslational modifications (PTMs) add “tubulin code” to confer function diversity of microtubules (He, Ling et al., 2020, Janke & Magiera, 2020). In quiescence cells, microtubule polyglutamylation occurs predominantly along the cilia axoneme and microtubules inside neuronal axons (He et al., 2020, Yang, Hong et al., 2021), with its physiological importance underappreciated until recently. Microtubule polyglutamylation is a reversible process coordinated by nine tubulin tyrosine ligase-like (TTLL) glutamylases (Janke, Rogowski et al., 2005, van Dijk, Rogowski et al., 2007) and six cytoplasmic carboxyl peptidase (CCP) deglutamylases (Rogowski, van Dijk et al., 2010). The physiological importance of polyglutamylation modification on different microtubule structures remain poorly understood. The activity of microtubule-severing enzyme spastin in neurons is stimulated by low-level polyglutamylation but inhibited gradually by increased polyglutamylation (Lacroix, van Dijk et al., 2010, Roll-Mecak & Vale, 2008, Valenstein & Roll-Mecak, 2016). Aberrant microtubule polyglutamylation in the brain may be associated with neural degeneration in human or mice (Rogowski et al., 2010, Shashi, Magiera et al., 2018).

Emerging evidence suggest that defective axoneme polyglutamylation disrupts cilia stability and function and leads to ciliopathies (He et al., 2020, He, Ma et al., 2018, Hong, Wang et al., 2018, Kanamaru, Neuner et al., 2022, Ki, Kim et al., 2020, Yang et al., 2021). However, how proper axoneme polyglutamylation is achieved remains poorly understood. Mutations in several ciliopathy Joubert Syndrome (JBTS) genes (*ARL13B*, *CEP41*, *KIF7, ARMC9* and *TOGARAM1*) cause defective axoneme polyglutamylation (He et al., 2018, He, Subramanian et al., 2014, Latour, Van De Weghe et al., 2020, Lee, Silhavy et al., 2012). The dominant mutations in Tau tubulin kinase 2 (TTBK2), a key ciliogenesis regulator, responsible for spinocerebellar ataxia type 11 (SCA11) also result in disrupted axoneme polyglutamylation (Bowie, Norris et al., 2018). We previously reported FIP5 (Rab11 Family Interacting Protein 5) as an interactor of JBTS protein ARL13B (He et al., 2018). ARL13B recruits FIP5 to cilia base upon ciliogenesis induction, and then FIP5-positive vehicles tether TTLL-positive vehicles to promote cilia import of TTLL5/6 glutamylases (He et al., 2018). Impaired axoneme polyglutamylation destabilizes cilia by increasing resorption, and importantly, damages cilia function by disrupting proper cilia localization of sensory receptors/molecules (He et al., 2018, Hong et al., 2018). Of note, enhancing axoneme polyglutamylation by depleting the primary cilia deglutamylase CCP5 strongly stabilizes cilia and enhances cilia function (He et al., 2018). The fact that *Ccp5^-/-^* mice are viable, fertile, and without ciliopathy-associated phenotypes supports the perspective that enhancing axoneme polyglutamylation is not detrimental to *in vivo* development and thus holds strong potential in future therapeutic development (Wu, Wei et al., 2017, Xia, Ye et al., 2016).

In this study, we uncover a core regulatory mechanism of axoneme polyglutamylation in which an uncanonical activity of CDK6, but not its close paralog CDK4, is activated by CDK7 and specifically suppresses axoneme polyglutamylation by phosphorylating FIP5 at the cilia base and inhibiting FIP5-dependent cilia import of tubulin glutamylases TTLL5/6. We further discovered that Polycystin-1 (PC1) deficiency leads to significantly upregulated CDK6 and impaired axoneme polyglutamylation in *PKD* cells. CDK6 inhibition, either genetically or pharmaceutically, strongly enhances axoneme polyglutamylation and restores defective cilia function in polyglutamylation-deficiency-associated ciliopathy models without adverse impacts on cell viability and health. We thus propose that targeting the ciliary CDK6-FIP5 pathway to enhance axoneme polyglutamylation holds promising potential to restore defective cilia signaling in ciliopathy treatment.

## Results

### CDK7 inhibition promotes axoneme polyglutamylation

Axoneme polyglutamylation has been known to be a dynamic regulation (Yang et al., 2021). Kinase mediated protein phosphorylation is a major regulatory mechanism for diverse cellular processes responding to signals arising inside or outside the cell. To identify key modulators in regulating axoneme polyglutamylation, we screened a collection of kinase inhibitors (Appendix Table S1) using glutamylated axoneme as a readout in hTERT-immortalized retinal pigment epithelial cells (RPE-1), a widely used model with consistent cilia length. Our preliminary screening revealed that the non-selective CDKs inhibitors (e.g., Flavopiridol hydrochloride, Milciclib, AZD5438) (Fig EV1), as well as the CDK7 specific inhibitors (THZ1, THZ2 and BS-181) (Fig 1A and EV1), could drastically and specifically enhance axoneme polyglutamylation in RPE-1 cilia, suggesting that CDK7 is a negative regulator of axoneme polyglutamylation.

**Figure 1.**
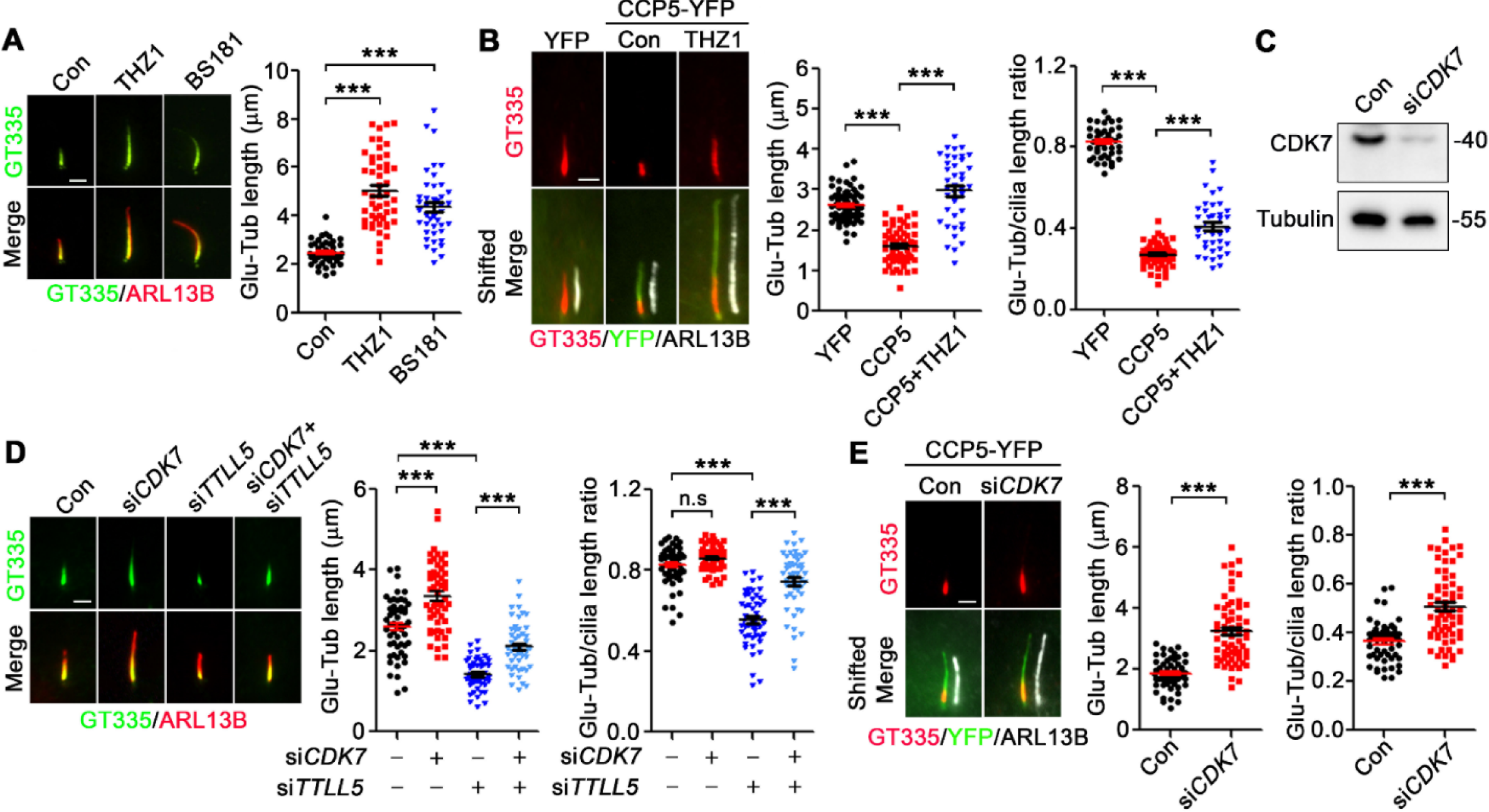
CDK7 suppresses axoneme polyglutamylation. **A** CDK7 selective inhibitors increase the length of glutamylated axoneme in RPE-1 cells. Cells were treated with THZ1 (1 mM) or BS-181 (10 mM) for 24 hrs in serum free medium. ARL13B was used as a marker of cilia. The glutamylated axoneme were labeled by antibody GT335. **B** THZ1 partially rescues the axoneme polyglutamylation in CCP5-YFP overexpressed RPE-1 cells. **C** The knockdown efficiency of CDK7 siRNA was accessed by western blotting. **D** Knockdown of CDK7 restores the axoneme polyglutamylation in TTLL5-depleted cells. **E** Knockdown of CDK7 partially rescues the axoneme polyglutamylation in CCP5-YFP overexpressed cells. Data information: Quantified data are presented as mean ± s.e.m. Statistical analyses were performed by one-way ANOVA analyses with Tukey’s post-hoc test for multiple comparisons (A, B, D) or two-tailed unpaired Student’s t-test (E). N≥40 (A) or 50 (B, D, E) cilia. *\*\*\*P* < 0.001. n.s: not significant. Scale bars: 2 μm.

Since increased axoneme polyglutamylation could be a secondary effect of cilia elongation, we examined if CDK inhibition directly promote axoneme polyglutamylation or cilia elongation. We used cell models with hypoglutamylated axoneme by knockdown of the ciliary glutamylase TTLL5 or overexpression of the ciliary deglutamylase CCP5 in RPE-1 cells. TTLL5 knockdown or CCP5 overexpression significantly impairs axoneme polyglutamylation, but not affect cilia length in RPE-1 cells (He et al., 2018, Hong et al., 2018). CDK7 specific inhibitor THZ1 effectively restores axoneme polyglutamylation and polyglutamylation-to-cilium ratio in CCP5-overexpressed cells (Fig 1B). Consistently, si*CDK7* treatment phenocopies THZ1 effect in WT, TTLL5-deficient, or CCP5-overexpressed RPE-1 cells (Fig 1C-E). These data suggest that CDK7 directly inhibits axoneme polyglutamylation. Notably, overexpression of TTLL5/6 leads to non-specific cytoplasmic polyglutamylation due to accumulation of glutamylases in the cytoplasm (He et al., 2018). Pharmacological or genetic inhibition of CDK7 did not lead to upregulated microtubule polyglutamylation in the cytoplasm (Fig EV2). This suggests that CDK7 specifically modulates the ciliary polyglutamylation but not affects cytoplasmic microtubules.

### A CDK7-CDK6 kinase cascade inhibits axoneme polyglutamylation

As expected, both overexpressed and endogenous CDK7 exclusively localize in the nucleus (Fig 2A). The absence of CDK7 in or around cilia suggests a missing link between the nuclear CDK7 activity and its specific regulation on axoneme polyglutamylation. CDK7 acts as the CAK (CDK activating kinase) by phosphorylating other CDKs, including CDK1, 2, 4, 6, and 9 (Fisher, 2005, Schachter, Merrick et al., 2013). We reasoned that it is likely one of the CDK7 substrates, e.g., downstream CDK(s), directly regulate axoneme polyglutamylation. By performing *siRNA* screening of CDKs activated by CDK7, we discovered that only knockdown of CDK6 significantly increased axoneme polyglutamylation (Fig 2B, C, and EV3). Notably, knockdown of CDK4, the close paralog of CDK6 in regulating cell-cycle transition (Malumbres, Sotillo et al., 2004), show no impact on axoneme polyglutamylation (Fig 2B and C). To further distinguish the role of CDK4 and CDK6 in regulation of axoneme polyglutamylation, we used CRISPR-Cas9 system in immortalized normal human renal cortical tubular epithelial (RCTE) cells to generate *CDK4^-/-^*and *CDK6^-/-^* cells (Fig 2D). Consistently, *CDK6^-/-^,* but not *CDK4^-/-^*, cells show significantly upregulated axoneme polyglutamylation (Fig 2E).

**Figure 2.**
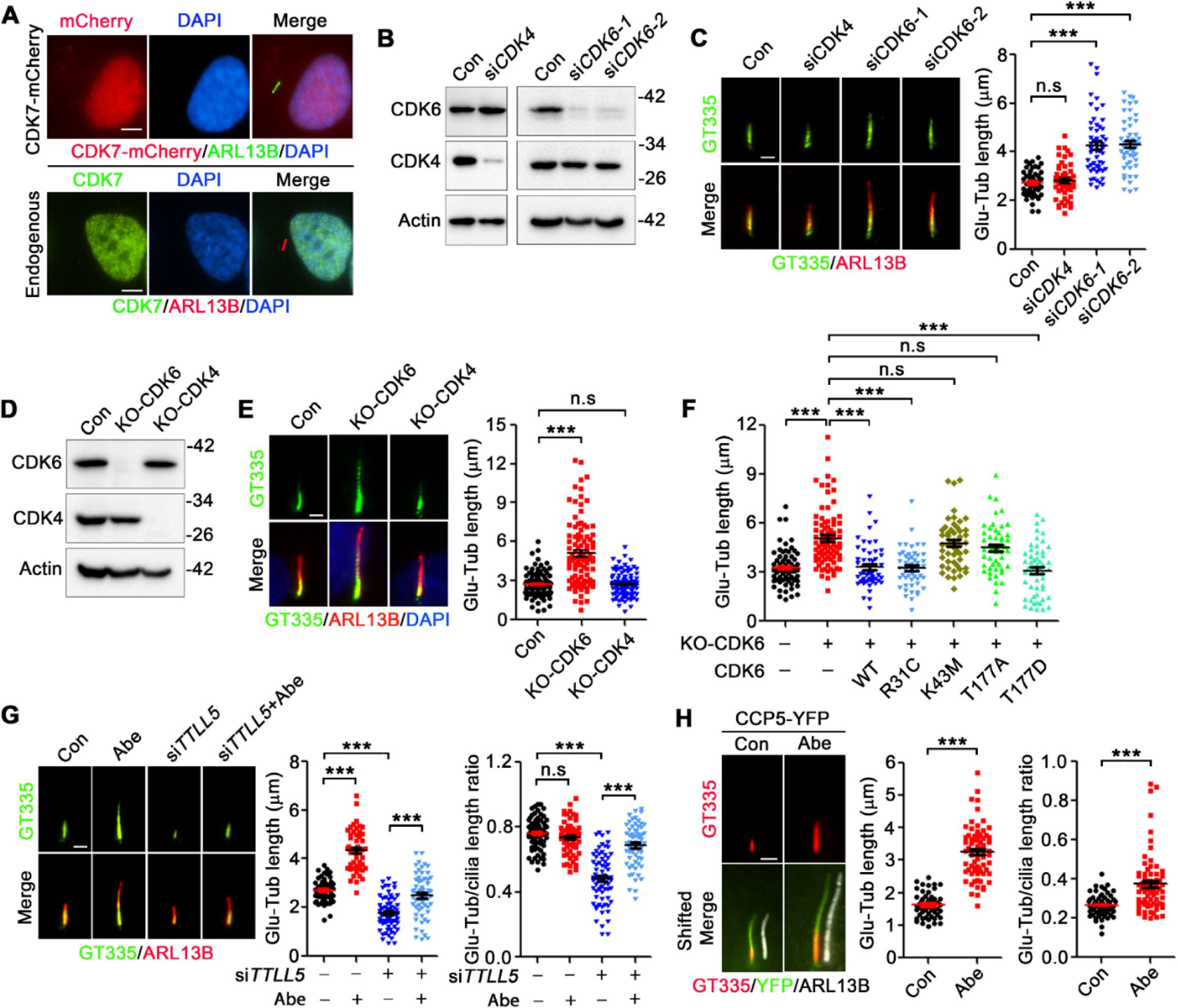
CDK6, but not CDK4, suppresses axoneme polyglutamylation. **A** Ectopically expressed and endogenous CDK7 localize at nucleus. **B** The knockdown efficiency of CDK4 and CDK6 specific siRNAs were accessed by western blotting. **C** Knockdown of CDK6, but not CDK4, promotes axoneme polyglutamylation in RPE-1 cells. **D** The knockout validation of CDK4 and CDK6 by western blotting in RCTE cells. **E** Knockout CDK6, but not CDK4, promotes axoneme polyglutamylation in RCTE cells. **F** The CDK6^-/-^ RCTE cells were transfected with indicated mCherry-tagged CDK6. 48 hrs later, cells were cultured in serum-free medium for 24 hrs to induce ciliogenesis. The length of polyglutamylated axoneme in mCherry-positive cells were measured. **G** Abemaciclib (Abe) restores axoneme polyglutamylation in TTLL5-depleted RPE-1 cells. Cells were transfected with TTLL5 siRNA for 48 hrs, followed by abemaciclib (500 nM) treatment for 24 hrs in serum free medium. **H** Abemaciclib partially rescues the axoneme polyglutamylation in CCP5-YFP overexpressed RPE-1 cells. Data information: Quantified data are presented as mean ± s.e.m. Statistical analyses were performed by one-way ANOVA analyses with Tukey’s post-hoc test for multiple comparisons (C, E, F, G) or two-tailed unpaired Student’s t-test (H). *\*\*\*P* < 0.001. n.s: not significant. N≥ 50 Cilia. Scale bars: 5 μm (A) or 2 μm (C, E, G, H).

Of note, re-expression of WT- or a hyperactive CDK6 variant (CDK6^R31C^), but not the kinase-inactive mutant (CDK6^K43M^), reduces axoneme polyglutamylation in *CDK6^-/-^* RCTE cells (Fig 2F), suggesting the kinase activity of CDK6 is essential for axoneme glutamylation. Abemaciclib and Ribociclib are CDK4/6 selective and safe inhibitors which have been approved by FDA for long-term breast cancer treatment (George, Qureshi et al., 2021). As expected, treating RPE-1 cells with Abemaciclib or Ribociclib greatly increased the length of glutamylated axoneme (Fig 2G and EV1). Consistently, Abemaciclib could also restore axoneme glutamylation in TTLL5-deficient or CCP5-overexpressed RPE-1 cells (Fig 2G and H). Similar to CDK7 inhibition, Abemaciclib treatment or CDK6 knockdown showed no impact on cytoplasmic tubulin glutamylation, suggesting its specificity in regulating cilia polyglutamylation (Fig EV2). It is known that CDK7 activates CDK6 by T177 phosphorylation during G1 phase progression (Kaldis, Russo et al., 1998, Schachter et al., 2013). To confirm the role of the CDK7-CDK6 kinase cascade in axoneme polyglutamylation, we generated and overexpressed CDK6^T177D^ (mimicking activated CDK6) and CDK6^T177A^ (mimicking inactive CDK6) in *CDK6^-/-^* RCTE cells. As expected, CDK6^T177D^, but not CDK6^T177A^, effectively reduced axoneme polyglutamylation (Fig 2F), further supporting the inhibitory role of the CDK7-CDK6 phosphorylation cascade in regulating axoneme polyglutamylation.

### CDK6 and its cyclin partner cyclin D3 enrich at the ciliary base

To understand how CDK6 regulates axoneme polyglutamylation, we carefully examined its subcellular localization in ciliated cells. Either endogenous or overexpressed CDK6, together with its cyclin partner cyclin D3, enrich at the ciliary base in RPE-1 cells (Fig 3A, C, and D). Intriguingly, overexpressed CDK4 also showed cilia base localization (Fig 3B), implying that CDK4 and CDK6 evolve together to gain cilia-related functions, while CDK6 specifically regulates axoneme polyglutamylation. To further confirm it is the ciliary CDK6 responsible for regulation of axoneme polyglutamylation, we took advantage of a validated cilia trapping system. By fusing with the C-terminal of subdistal appendage protein CEP170 (CEP170C) (Guarguaglini, Duncan et al., 2005), protein of interest can be specifically trapped to the distal end of the basal body (Fig 3E-G). CDKs alone usually exhibit weak kinase activity, while the cyclin-CDK fusion proteins are constitutively active (Rao, Stamm et al., 1999). We thus constructed the Cyclin D3-CDK6 fusion protein (D3K6) as described previously (Fig 3E) (Rao et al., 1999). Remarkably, the cilia-trapped CEP170C-D3K6-YFP disrupted axoneme polyglutamylation in RCTE or RPE-1 cells (Fig 3F and G), demonstrating the functional site for CDK6-regulated axoneme polyglutamylation is the ciliary base.

**Figure 3.**
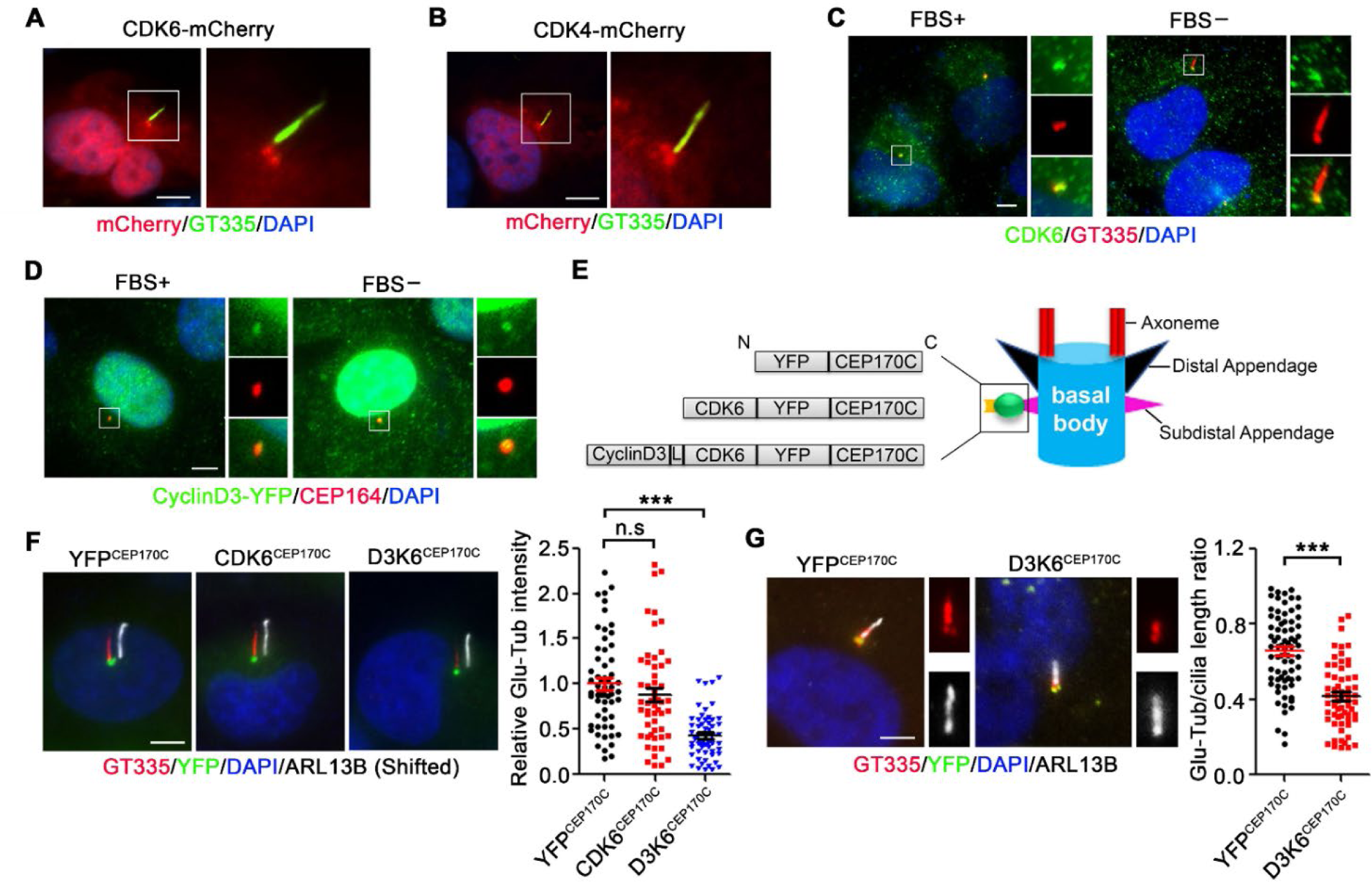
Ciliary CDK6 is sufficient to suppress axoneme polyglutamylation. **A, B** The subcellular localization of ectopically expressed CDK6-mCherry (A) and CDK4-mCherry (B) in ciliated RPE-1 cells. **C** The subcellular localization of endogenous CDK6 in non-ciliated (FBS+) and ciliated (FBS-) RPE-1 cells. **D** The subcellular localization of ectopically expressed cyclin D3-YFP in indicated RPE-1 cells. CEP164 were used as the marker of mother centriole or basal body. **E** Schematic illustration of CEP170C-based cilia targeting system. N: N-terminal; C: C-terminal; L: flexible linker. **F, G** Ciliary targeting of CyclinD3-CDK6 inhibits axoneme polygultamylation. The mean fluorescent intensity of glutamylated axoneme in indicated YFP-positive RCTE cells (F) or the ratio of glutamylated axoneme length to cilia length in indicated YFP-positive RPE-1 cells (G) were quantified. Data information: Quantified data are presented as mean ± s.e.m. Statistical analyses were performed by one-way ANOVA analyses with Tukey’s post-hoc test for multiple comparisons (F) or two-tailed unpaired Student’s t-test (G). *\*\*\*P* < 0.001. n.s: not significant. Scale bars: 5 μm. N≥50 Cilia.

### The CDK7-CDK6 cascade suppresses cilia import of tubulin glutamylases

We previously discovered that tubulin glutamylases TTLL5 and TTLL6 are responsible for axoneme polyglutamylation (He et al., 2018). TTLL5/6 exhibit dynamic cilia localization patterns (He et al., 2018), suggestive of an actively regulated cilia import of glutamylases. To test if the CDK7-CDK6 kinase cascade regulates the ciliary import of tubulin glutamylases, we examined TTLL5/6 localization in CDK7- or CDK6-deficient cells (Fig 4). Immediately after THZ1 or Abemaciclib treatment (2 hours after administrating drugs) (Fig 4A and B), or knockdown of *CDK6* or *CDK7* (Fig 4C-F), profoundly upregulated the ciliary level of TTLL5/6, suggesting that the CDK7-CDK6 cascade acts as the rate-limiting factor in control of cilia import of glutamylases.

**Figure 4.**
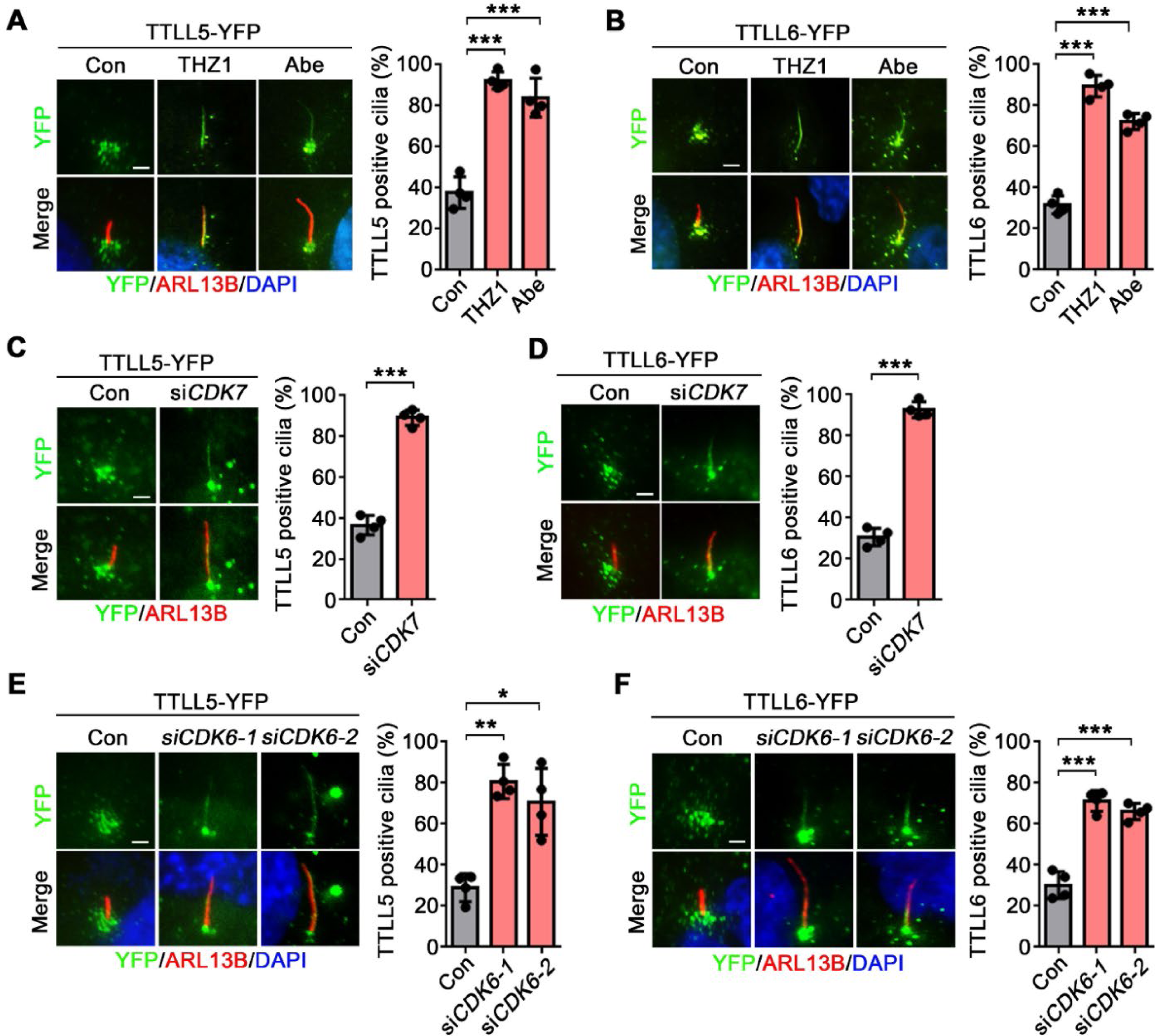
CDK7 and CDK6 suppress ciliary import of TTLL5 and TTLL6. **A, B** THZ1 and Abemaciclib promotes ciliary import of TTLL5 (A) and TTLL6 (B). After 24 hrs serum starvation, the TTLL5-YFP or TTLL6-YFP overexpressed RPE-1 cells were treated with THZ1 or Abemaciclib for 2 hrs. **C, D** Knockdown of CDK7 promotes ciliary import of TTLL5 (C) and TTLL6 (D). **E, F** Knockdown of CDK6 promotes ciliary import of TTLL5 (E) and TTLL6 (F). Data information: Quantified data are presented as mean ± s.d. Statistical analyses were performed by one-way ANOVA analyses with Tukey’s post-hoc test for multiple comparisons (A, B, E, F) or two-tailed unpaired Student’s t-test (C, D). N=4 independent experiments. *\*P* < 0.05; *\*\*P* < 0.01; *\*\*\*P* < 0.001. Scale bars: 2 μm.

### CDK6 inhibits axoneme polyglutamylation by suppressing the ciliary accumulation of FIP5

The RAB11 family, which includes RAB11A, RAB11B and RAB25, is the major GTPase regulators of vesicular trafficking from endosomes to other cellular membranes (Kelly., Horgan. et al., 2012). RAB11A is the predominant one that ubiquitously expresses in most cell types. We previously found that FIP5, an effector of RAB11, is recruited to the proximity of the basal body to ensure the ciliary import of TTLL5/6 shortly after serum starvation-induced ciliogenesis (He et al., 2018). Of note, Abemaciclib treatment strongly recruits FIP5 to the proximity of the centrosome, even without serum starvation (Fig 5A; Movie EV1). Moreover, cilia-trapped CEP170C-D3K6-YFP effectively blocked the ciliary recruitment of FIP5 (Fig 5B). We confirmed the loss of the ciliary FIP5 is not due to the change of its protein stability (Appendix Fig S1). Consistently, Abemaciclib treatment, or *CDK6* ablation, induced axoneme hyperglutamylation (Fig 5C-E) and cilia import of TTLL5/6 (Fig 5F and G) were suppressed by FIP5 depletion. Collectively, these data suggest that activated CDK6 at the ciliary base suppresses axoneme polyglutamylation by inhibiting the ciliary accumulation of FIP5, and thus blocking FIP5-dependent cilia import of tubulin glutamylases.

**Figure 5.**
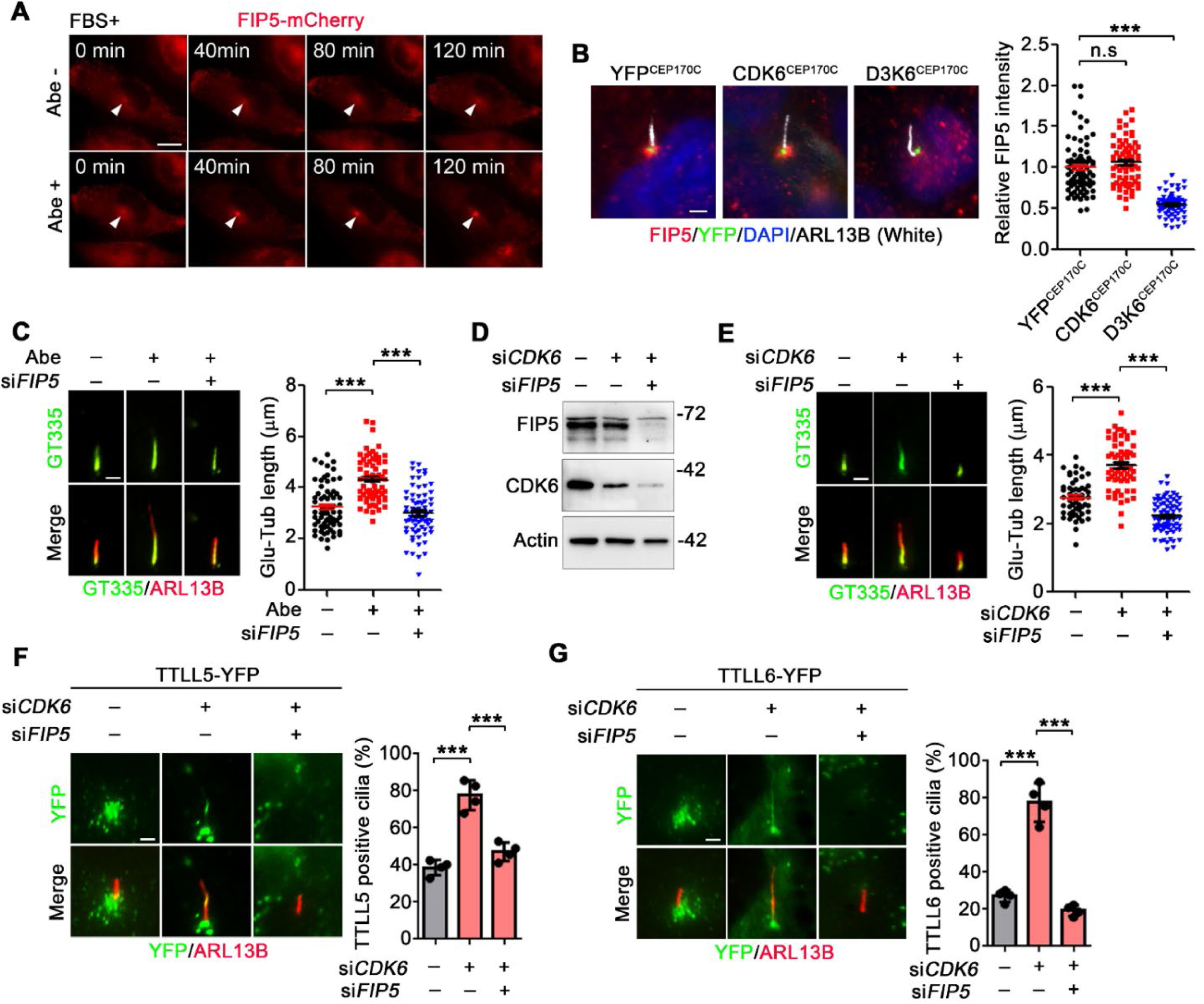
FIP5 acts downstream of CDK6 to regulate axoneme polyglutamylation. **A** The FIP5-mCherry overexpressed RCTE cells were cultured in serum-containing medium. Images cropped from live-cell imaging before and after Abemaciclib treatment. White triangles indicate the accumulation loci of FIP5-mCherry-positive vesicles. **B** The subcellular localization, and fluorescent intensity of FIP5 at cilia base in indicated YFP-positive RCTE cells. **C** Knockdown of FIP5 inhibits Abemaciclib induced axoneme hyperglutamylation in RPE-1 cells. **D** The knockdown efficiency of CDK6 and FIP5 siRNAs were accessed by western blotting. **E** Knockdown of FIP5 inhibits CDK6 depletion induced axoneme hyperglutamylation in RPE-1 cells. **F, G** Knockdown of FIP5 inhibits CDK6 depletion induced ciliary import of TTLL5 (F) and TTLL6 (G). Data information: Quantified data are presented as mean ± s.e.m. (B, C, E) or s.d. (F, G). Statistical analyses were performed by one-way ANOVA analyses with Tukey’s post-hoc test for multiple comparisons. N≥ 50 Cilia (B, C, E) or =4 independent experiments (F, G). *\*\*\*P* < 0.001. n.s: not significant. Scale bars: 10 μm (A), 5 μm (B) or 2 μm (C, E, F, G).

### CDK6 phosphorylates FIP5 at S641 to suppress FIP5 accumulation at the ciliary base

CDKs use cyclins to recognize substrates and then phosphorylate at CDK consensus site(s) (full sequence: [S/T*]PX[K/R] (Ord, Moll et al., 2019)). Of note, FIP5 has six cyclin-binding motifs (RXL) and two strong and six weak CDK consensus sites (Fig 6A), raising an intriguing hypothesis that FIP5 may be a direct substrate of CDK6. Indeed, by using anti-phosphoserine antibody, strong phosphorylation of FIP5 could be detected only when co-expressed with D3K6 in 293T cells (Fig 6B). We then assessed CDK6 phosphorylation site(s) on FIP5 by performing site-directed mutagenesis. For all tested phospho-silent (S-A) mutants, only overexpression of FIP5^S641A^ induced axoneme hyperglutamylation in RCTE cells (Fig 6C). FIP5 phosphorylation was markedly reduced when co-expressing FIP5^S641A^ with D3K6 in 293T cells (Fig 6D), suggesting that FIP5 S641 is a genuine CDK6 phosphorylation site. In contrast to WT-FIP5, FIP5^S641A^ exhibited constitutive centrosome/cilia accumulation without serum starvation, reminiscent of the phenotype of Abemaciclib-treated cells (Fig 6E). Consistently, overexpression of cilia-trapped D3K6 disrupted the accumulation of FIP5, but not FIP5^S641A^, at the ciliary base (Fig 6F). Collectively, these data demonstrate that CDK6 suppresses FIP5 accumulation at the ciliary base by phosphorylated FIP5-S641.

**Figure 6.**
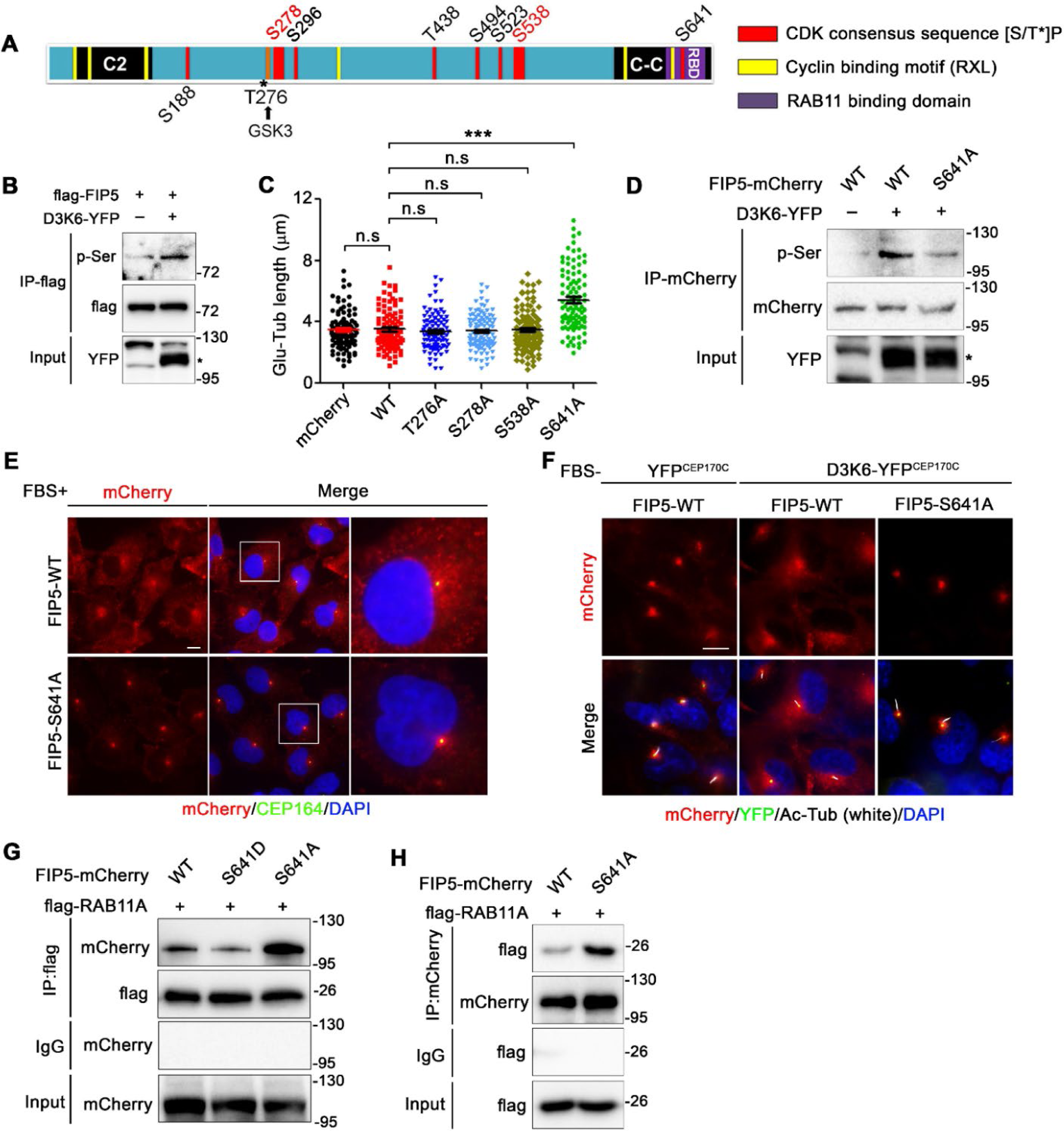
S641 phosphorylation by CDK6 interrupts cilia base accumulation and RAB11 interaction of FIP5. **A** The putative CDK6 phosphorylation sites and cyclin binding motifs in FIP5. **B** flag-tagged FIP5 and CyclinD3-CDK6-YFP were transfected in 293T cells. The phosphorylation of FIP5 were examined by anti-phosphorylated Serine antibody after immunoprecipitation of flag-FIP5. * indicates the positive band of D3K6-YFP. **C** mCherry or mCherry-tagged FIP5 mutants were transfected in RCTE cells. The length of glutamylated axoneme in mCherry-positive cells were measured. **D** Indicated mCherry-tagged FIP5 and CyclinD3-CDK6-YFP were transfected in 293T cells. The phosphorylation of FIP5 were examined by anti-phosphorylated Serine antibody after immunoprecipitation of FIP5-mCherry. **E** Subcellular localization of mCherry-tagged WT- or S641A-FIP5 in non-ciliated RCTE cells. The enlarged fields indicated by white squares and showed in the right panels. **F** The subcellular localization of mCherry-tagged FIP5 in indicated YFP-positive ciliated RCTE cells. **G, H** Indicated mCherry-tagged FIP5 and flag-tagged RAB11A were transfected in 293T cells. The interaction between FIP5 and RAB11A were examined by co-immunoprecipitation. Data information: Quantified data are presented as mean ± s.e.m. Statistical analyses were performed by one-way ANOVA analyses with Tukey’s post-hoc test for multiple comparisons. N≥100 Cilia. *\*\*\*P* < 0.001. n.s: not significant. Scale bars: 10 μm.

Besides S641 phosphorylation, FIP5-T276 phosphorylation by GSK-3 regulates apical trafficking of endosomes during mitosis (Li, Mangan et al., 2014). FIP5-S188 phosphorylation by ERK controls the transcytosis of the polymeric immunoglobulin receptor (Su, Bryant et al., 2010). We show that overexpression of FIP5^T276A^ or treatment of ERK inhibitors did not increase axoneme polyglutamylation (Fig 6C and EV1), suggesting the specificity of FIP5-S641 phosphorylation in regulating axoneme polyglutamylation.

### Phosphorylation of FIP5 at S641 disrupts RAB11-FIP5 interaction

During ciliogenesis, RAB11 is recruited to the ciliary base by FIP5, and RAB11-FIP5 complex is essential for the ciliary import of glutamylases (He et al., 2018). Intriguingly, S641 site localizes within the RAB11-binding domain (RBD) of FIP5 (Fig 6A), implying S641 phosphorylation may influence FIP5/RAB11 interaction. By co-expressing RAB11A and FIP5 variants in 293T cells, we discovered that FIP5^S641A^ exhibits significantly enhanced interaction with RAB11A (Fig 6G and H). As expected, S641D (phospho-mimetic) mutation disrupted association between FIP5 and RAB11A (Fig 6G). We concluded that CDK6-mediated S641 phosphorylation suppress axoneme polyglutamylation by disrupting RAB11-FIP5 interaction, and thus, prevents proper ciliary trafficking of TTLL5/6.

### Abemaciclib restores polyglutamylation and ciliary signaling in Joubert syndrome cells

Polyglutamylation deficiency has been associated with mutations of multiple JBTS genes (He et al., 2018, He et al., 2014, Latour et al., 2020, Lee et al., 2012). We and others show that genetic ablation of the ciliary deglutamylase CCP5 can restore defective axoneme polyglutamylation caused by deficiency of TTLL5/6 or JBTS gene *CEP41* (He et al., 2018, Ki et al., 2020). We next explored whether CDK6 inhibition could be used to enhance axoneme polyglutamylation and restore cilia function in polyglutamylation deficiency-associated JBTS cells. NIH-3T3 cells are commonly used for studying ciliary Sonic hedgehog (*Shh*) signaling and the mouse inner medullary collecting duct cells (IMCD3) exhibit robust ciliary localization of Polycystin-2 (PC2). We first confirmed that *Cep41* and *Armc9* knockdown with validated siRNA (Lee et al., 2012) significantly reduced axoneme polyglutamylation in NIH-3T3 (Fig 7A and B, and Appendix Fig S2A) and IMCD3 cells (Fig 7E and F, and Appendix Fig S2B). Remarkably, Abemaciclib treatment efficiently restored the axoneme polyglutamylation in *siCep41* and *siArmc9* cells (Fig 7A, B, E, and F). In agreement with the critical role of axoneme polyglutamylation in *Shh* signaling and Polycystin signaling (He et al., 2018, Hong et al., 2018), *siCep41* and *siArmc9* cells show aberrant *Shh* signaling including impaired cilia tip localization of GLI3 (Fig 7A and C), suppressed *Gli1*/*Ptch1* transcription upon *Shh* agonist SAG stimulation (Fig 7D), and compromised cilia localization of PC2 (Fig 7E and G). Abemaciclib treatment effectively corrects *Shh* signaling (Fig 7A-D) and restores the ciliary level of PC2 (Fig 7E-G) in *siCep41* and *siArmc9* cells. These data suggest that inhibition of CDK6 restores cilia function in JBTS cells with defective axoneme polyglutamylation.

**Figure 7.**
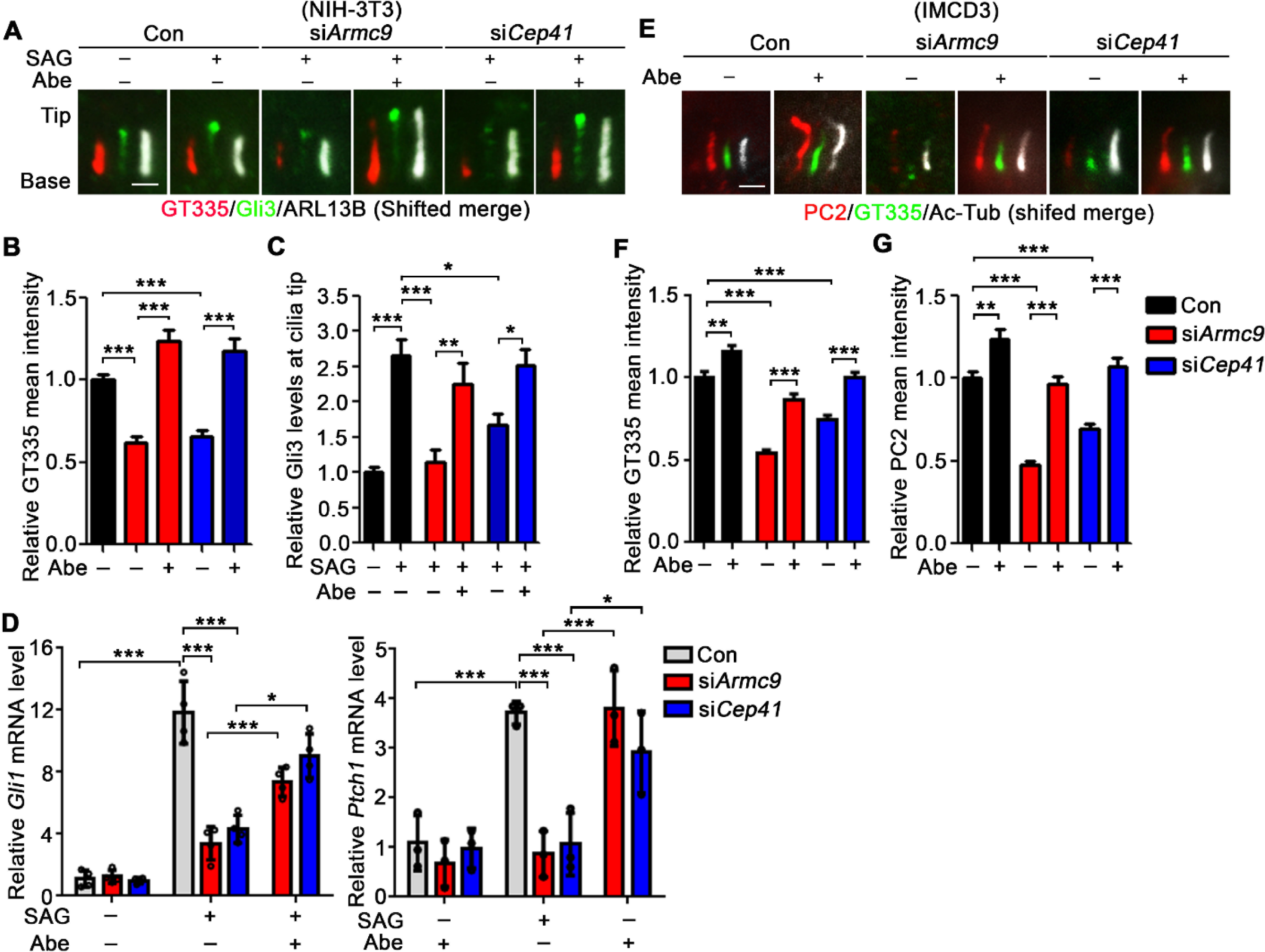
Abemaciclib rescues glutamylation and ciliary signaling in JBTS mutant cells. **A-D** Abemaciclib (200 nM, 24 hrs) restores axoneme polyglutamylation and Shh signaling in *Armc9* and *Cep41* knocked-down NIH-3T3 cells, examined by mean intensity of GT335 in cilia (A and B), SAG (100 nM, 24 hrs) induced cilia tip accumulation of GLI3 (A and C), and Gli1 and Ptch1 transcription (D). **E**-**G**. Abemaciclib (200 nM, 24 hrs) restores axoneme polyglutamylation (E and F) and ciliary localization of PC2 (E and G) in *Armc9* and *Cep4*1 knocked-down IMCD3 cells. Data information: Quantified data are presented as mean ± s.e.m. (B, C, F, G) or s.d. (D). Statistical analyses were performed by one-way ANOVA analyses with Tukey’s post-hoc test for multiple comparisons. N≥100 Cilia (B, C, F, G) or =3 or 4 independent experiments (D). *\*P* < 0.05; *\*\*P* < 0.01; *\*\*\*P* < 0.001. Scale bars: 2 μm.

### CDK6 upregulation and impaired axoneme hypoglutamylation are implicated in ADPKD

ADPKD, the most common life-threatening genetic disease, is mainly caused by mutations in *PKD1* or *PKD2*, which encodes PC1 and PC2, respectively (Torres & Harris, 2006). Recent studies suggest that functional polycystin dosage governs ADPKD disease severity (Dong, Zhang et al., 2021, Hopp, Ward et al., 2012, Lakhia, Ramalingam et al., 2022, S., VJ. et al., 2009). Remarkably, increasing the expression of PC2 alone can slow cyst growth even in *Pkd1*-mutant mice (Lakhia et al., 2022). As axoneme polygultamylation controls the ciliary abundance of polycystins, we investigated the involvement of CDK6-FIP5 pathway in ADPKD. Intriguingly, abnormal upregulation of CDK6, but not the other CDKs, was observed in immortalized human ADPKD renal cyst-derived *PKD1^-/+^* cells (Fig EV4A), as well as in renal tubules of ADPKD patients or hypomorphic *Pkd1*^RC/RC^ mice (Hopp et al., 2012), particularly in cyst-lining cells (Fig 8A and B). Consistent with the inhibitory role of CDK6 on FIP5, both immortalized human *PKD1^-/+^*cells (Fig EV4B) and primary mouse *Pkd1^RC/RC^* renal tubular epithelial cells (Fig 8C) exhibited a significant reduction in axoneme polyglutamylation. Reminiscent of the effects observed in JBTS cells, treatment of Abemaciclib effectively restored axoneme polyglutamylation in ADPKD cells (Fig 8C and EV4B). To explore the potential involvement of defective axoneme polyglutamylation in renal cystogenesis in ADPKD, we employed an *ex vivo* metanephric cystogenesis model using embryonic kidneys from *Pkd1^RC/RC^* mice. Remarkably, Abemaciclib treatment strongly suppressed the cystogenesis in *Pkd1^RC/RC^* embryonic kidneys, even at a low dosage of 50 nM, without adversely affecting kidney growth (Fig 8D). These data collectively highlight that CDK6 as a promising therapeutic target for restoring defective cilia function in ciliopathies associated with polyglutamylation deficiency, including ADPKD.

**Figure 8.**
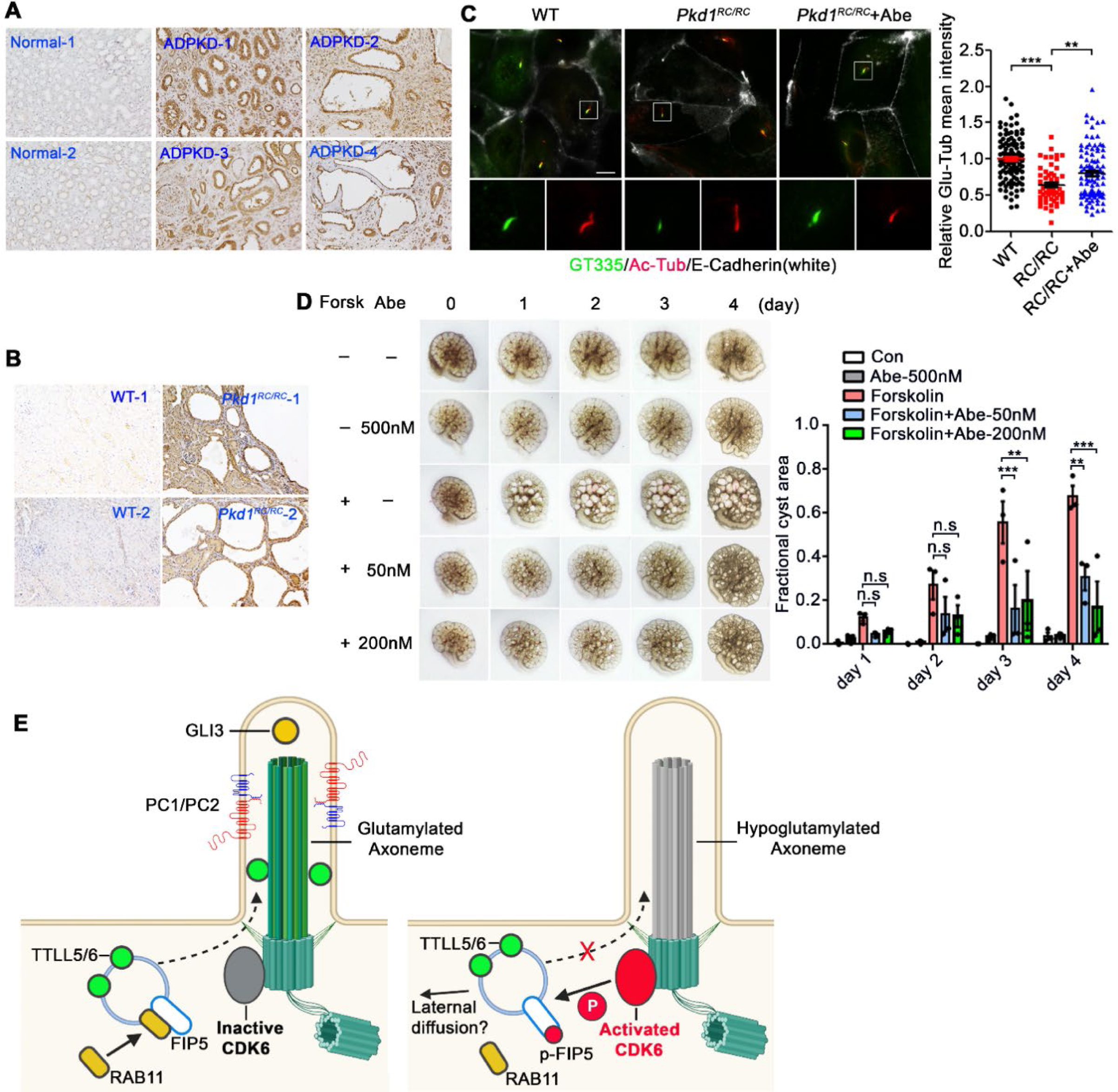
Upregulation of CDK6 and defective axoneme polyglutamylation are implicated in ADPKD. **A, B** Abnormal upregulation of CDK6 in renal tubules of ADPKD patients (A) and *Pkd1^RC/RC^* mice (B) as shown by immunohistochemistry. **C** Primary renal epithelial cells isolated from WT and *Pkd1^RC/RC^* mouse at 1-year-old were used to assess the level of axoneme polyglutamylation by IF. E-Cadherin was used as a marker of epithelial cells. The mean intensity of GT335 in cilia of E-Cadherin positive cells were quantified. **D** E13.5 embryonic kidneys from *Pkd1^RC/RC^* mice were cultured and treated with Forskolin (Forsk, 10 μM) or Abemaciclib as indicated. Representative pictures of kidneys and quantification of cyst area at indicated time points are shown. **E** Schematic model illustrating the CDK6-FIP5 phosphorylation cascade-regulated axoneme polyglutamylation. In cells with inactivated CDK6, FIP5 interacts with RAB11 to promote the ciliary import of TTLL5/6 and subsequent axoneme polyglutamylation. Glutamylated axoneme supports the cilia localization of polycystins and signaling molecules such as Hedgehog component GLI3. In cells with activated CDK6, CDK6 at the cilia base phosphorylates FIP5 at S641, disrupting the FIP5-RAB11 interaction and subsequent ciliary import of TTLL5/6. This leads to axoneme hypoglutamylation and defective ciliary localization of polycystins and GLI3. Image created with BioRender.com. Data information: Quantified data in C are presented as mean ± s.e.m. Statistical analyses were performed by One-way ANOVA with Tukey’s post-hoc test for multiple comparisons. N≥55 Cilia. Quantified data in D are presented as mean ± s.e.m. Statistical analyses were performed by Two-way ANOVA with multiple comparisons. N=3 embryonic kidneys. n.s: not significant; *\*\*P* < 0.01; *\*\*\*P*<0.001. Scale bar: 2 μm.

## Discussion

In summary, our study uncovers a novel CDK7-CDK6-FIP5 phosphorylation cascade in the control of cilia import of tubulin glutamylases and thereby cilia signaling (e.g., Polycystin, Hedgehog signaling) (Fig 8E). We and others demonstrated that axoneme polyglutamylation stabilizes cilia by modulating cilia resorption rate (Bowie et al., 2018, He et al., 2018, Hong et al., 2018, Kanamaru et al., 2022, Ki et al., 2020). During cell cycle progression, CDK7 activates CDK4/6 to promote G1 progression (Kaldis et al., 1998, Schachter et al., 2013). Cilia stability has been known to influence cell cycle progression (Kasahara & Inagaki, 2021), but with molecular links remaining poorly understood. Our current discovery that the CDK7-CDK6-phosphorylation cascade suppresses axoneme polyglutamylation suggests an intriguing regulation in which activation of this cascade during cell cycle not only promotes G0-to-G1 transition, but also destabilizes the cilium so that the basal body can be transformed back to the mother centriole to allow cell cycle re-entry. On the other hand, inactivation of this cascade during cell cycle exit naturally promotes axoneme polyglutamylation, enhances cilia stability, and thus supports cilia function in quiescent cells. Constitutive localization of CDK6 both in the nucleus and at the ciliary base suggest that activated CDK6 may be constantly cycled between the nucleus and the ciliary base in ciliated cells.

CDK4 and CDK6 are paralogs with redundant function in G1 phase of the cell cycle (Kozar & Sicinski, 2005, Malumbres et al., 2004). Of note, both CDK4 and CDK6 are found at the basal body (Fig. 3 and (Li, Zhou et al., 2020)), but with clearly different functions. Function heterogeneity for CDK4 and CDK6 has been documented in a variety of biological scenarios (Grossel & Hinds, 2006, Wang, Nicolay et al., 2017). It is not surprising that only CDK6, but not CDK4, suppresses axoneme polyglutamylation. Not like *CDK4^-/-^* mice that are born with small size, hypoplasia of various organs, and infertility, *CDK6^-/-^* mice are viable, fertile, and anatomically normal, indicative of complete compensation of CDK6 function by CDK4 during cell cycle progression but not *vice versa* (Kozar & Sicinski, 2005). Thus, the specificity of CDK6 in regulation of axoneme polyglutamylation makes CDK6-specific inhibition a promising strategy in future development of ciliopathy therapeutics.

Phosphorylation has been implicated in regulating FIP5 subcellular localization (Li et al., 2014, Su et al., 2010). Notably, CDK6 inhibition immediately triggers the accumulation of FIP5-positive vesicles at the ciliary base (Movie S1), suggestive of a fast change of endocytic trafficking. Kinesin-2, the microtubule plus end-directed molecular motor, interacts with the C-terminal of FIP5 to drive endosome transport during epithelial lumen formation (Li et al., 2014, Schonteich., Wilson. et al., 2008). It remains intriguing to investigate if a similar mechanism may be involved in cilia trafficking of FIP5-vesicles, which could be regulated by FIP5-S641 phosphorylation and FIP5-RAB11 interaction. Note that the basal body acts as the microtubule-organizing center (MTOC) of the minus-end of microtubule (O’Connell, 2021), the motor involved may belong to dynein family or minus-end-directed kinesins.

The physiological importance of axoneme glutamylation have been overlooked until recent correlation between disrupted axoneme polyglutamylation and human ciliopathies was identified (Bowie et al., 2018, He et al., 2018, He et al., 2014, Latour et al., 2020, Lee et al., 2012). Intriguingly, we observed significant CDK6 upregulation and impaired axoneme polyglutamylation in ADPKD. Prior studies, along with our current data, suggest that CDK6 inhibition generally show no adverse impacts on cell proliferation (Appendix Fig S3), probably due to the redundant role of CDK4 in cell cycle regulation (26). Intriguingly, in the context of ADPKD, a disease that associated with hyperproliferation of renal tubular cells, it is noteworthy that CDK6, but not other CDKs crucial for cell cycle progression, is upregulated in *PKD* cells. This observation suggests that the role of CDK6 in ADPKD is independent of cell proliferation but specifically associated with cilia dysfunction related to axoneme hypoglutamylation. One important question remains to be addressed is how impaired polycystin signaling leads to upregulated expression of CDK6.

Considering enzyme-regulated axoneme polyglutamylation is a druggable pathway, pharmacological restoration of polyglutamylation deficiency could be a promising therapeutic strategy in treatment of glutamylation-deficiency-associated ciliopathy. However, this approach is currently impeded by lack of specific inhibitors or agonists for cilia glutamylases or deglutamylases, respectively. Excitingly, our discovery that inhibition of CDK6-mediated FIP5 phosphorylation effectively rescues axoneme polyglutamylation and cilia signalings in glutamylation*-*deficient JBTS and ADPKD cells and suppresses *ex vivo* renal cystogenesis highlights the therapeutic potential of targeting CDK6. Further, CDK6 inhibition specifically enhances axoneme polyglutamylation but not global microtubule glutamylation, avoiding potential detrimental impacts on cytoplasmic microtubules. This is also supported by existing clinic evidence that CDK6 inhibitors show favorable safety profile in human (George et al., 2021). We also speculate that targeting the CDK6-FIP5 pathway may be applied in broader scenarios where enhancement of cilia function can be beneficial under either pathological or normal condition.

## Materials and Methods

### Cells and constructs

Cell lines used in this study were obtained from American Type Culture Collection. hTERT-RPE-1, RCTE, *PKD1*^-/+^ ADPKD patient-derived renal tubular cells (shared by Dr. Peter Harris) and IMCD3 cells were cultured in DMEM/F12 (Gibco, 11320033), containing 10% fetal bovine serum, and supplemented with penicillin and streptomycin. HEK-293T and NIH-3T3 cells were cultured in DMEM (Gibco, 10566016), containing 10% fetal bovine serum, and supplemented with penicillin and streptomycin. All the cells were cultured in a humidified atmosphere of 5% CO_2_ at 37°C.

The DNA template of CEP170C is a gift from Dr. Takanari Inoue (Stanford University School of Medicine). FIP5, CDK6, CDK4 and CDK7 are tagged with mCherry at the C-terminus and cloned into vector pCDH-CMV-MCS-EF1-Neo. TTLL5/6-YFP, CCP5-YFP, CyclinD3-L-CDK6-YFP, YFP-CEP170C, CDK6-YFP-CEP170C and CyclinD3-L-CDK6-YFP-CEP170C were cloned into vector pCDH-CMV-MCS-EF1-Neo as well. Flag-FIP5 was cloned into vector pcDNA3.1 (-). Site-specific mutagenesis of double-stranded plasmid DNAs were constructed using the Q5 Site-Directed Mutagenesis Kit (E0554S, New England Biolabs), according to the manufacturer’s instructions.

Stable cell lines were constructed using lentivirus system. For producing lentiviral particles, the pCDH constructs, together with psPAX2 (Addgene #12260) and pMD2.G (Addgene #12259), were transfected into HEK293T cells in the ratio of 4:3:1. Medium was collected 48 and 72 h after transfection. Lentiviral particles were further concentrated using Lenti-X Concentrator (Takara, 631231). After that, target cells were infected with lentivirus overnight. At 48 h after virus infection, the infected cells were further selected by G418 for 3–7 days.

### Kinase inhibitors screen

The kinase inhibitors inhibitor library was purchased form TargetMol (L1600, TargetMol). For drug screen, hTERT-RPE-1 cells were cultured in 24-well plate to reach confluence and then further serum starved and treated with kinase inhibitors (1 mM) or DMSO control for 24 hrs. After that, the cells were subjected to PFA fixation and conventional immunofluorescent assays. Primary cilia were identified using cilia marker ARL13B and the length of glutamylated axoneme were measured.

### Plasmids and siRNAs transfection

For plasmid transfection, X-tremeGENE 9 (Millipore Sigma, 6365779001) was used, according to the manufacturer’s instructions. siRNA duplexes were obtained from Invitrogene, and RNAi negative control were purchased from GE Healthcare Dharmacon. For siRNAs transfection, Lipofectamine RNAiMAX (Invitrogene; 13778-075) was used, according to the manufacturer’s instructions. Sequences of siRNA targeting corresponding mRNAs are as follows:

si*CDK1*: 5′-GAUCAACUCUUCAGGAUUU-3′ si*CDK2*: 5′-GCCAGAAACAAGUUGACGG-3′ si*CDK4*: 5′-AACAUUCUGGUGACAAGUGGU-3′

si*CDK5*: 5′-ACCUACGGAACUGUGUUCAAGGCCA-3′ si*CDK6-1*: 5′-AACAGACAGAGAAACCAAACU-3′ si*CDK6-2*: 5′-AACGUGGUCAGGUUGUUUGAU-3′ si*CDK7*: 5′-CCGCCUUAAGAGAGAUAAA-3′

si*CDK9*: 5′-GGUGCUGAUGGAAAACGAG-3′

si*FIP5*: 5′-CCAAGGUCUCCCUUCAGCAAGAUCA-3′ si*TTLL5*: 5′-AAGAACUCUUCCAGGAUUCUU-3′ si*Cep41*: 5′-UAGACAAAGGGCUCGUAAA-3′ si*Armc9*: 5′-UAGACAAAGGGCUCGUAAA-3′

### Gene deletion using the CRISPR–Cas9 system

CDK6-KO and CDK4-KO RCTE cells were generated through the CRISPR-Cas9 genomic editing system. Cells were transiently transfected with Cas9 and single guide RNAs plasmids with EGFP expression (PX458; Addgene plasmid no. 48138). The single guide RNAs targeting CDK6 and CDK4 (sequences TTAGATCGCGATGCACTACT and ATCTCGGTGAACGATGCAAT respectively) were predicted by the sgRNA Designer (https://portals.broadinstitute.org/gpp/public/analysis-tools/sgrna-design). Two days after transfection, individual cells expressing EGFP were sorted into 96-well plates through FACS (BD, FACSAria III). The KO clones were validated by western-blot analysis.

### Western blotting

For preparing the cell samples, cells were washed three times with PBS and lysed by 1×SDS loading buffer on ice and sonicated for 10 s. After boiling for 10 min at 95°C, protein samples were subjected to standard SDS-PAGE and transferred to polyvinylidene fluoride membranes. The membranes were blocked in 2% BSA for 1 h and incubated overnight at 4 °C with primary antibodies. After washing with TBS-T (Tris-buffered saline, 0.05% TWEEN) three times for 10 min each, the membranes were incubated with secondary antibodies for 1 hr at room temperature. After washing with TBS-T three times for 10 min each, the membranes were developed with chemical luminescence (BIO-RAD). Images were obtained using ChemiDoc Touch Imaging System (BIO-RAD). The following commercially available antibodies were used for western blot: RAB11FIP5 (14594-1-AP, Proteintech, dilution 1:1000), GFP (50430-2-AP, Proteintech, dilution 1:1000), Flag (8146, Cell Signaling Technology, dilution 1:2000), RAB11A (71-5300; Invitrogen, dilution 1:1000), anti-Phosphoserine (P5747, Sigma-Aldrich, dilution 1:500), CDKs (CDK Antibody Sampler Kit #9868; Cell Signaling Technology, dilution 1:1000), mCherry (16D7, Invitrogen, dilution 1:1000). β-Actin (3700, Cell Signaling, dilution 1:5000), α-Tubulin (T9026, Sigma-Aldrich, 1:5000).

### Immunofluorescence and live-cell imaging

For most of the staining, cells were fixed in 4% paraformaldehyde for 15 min at room temperature. For staining of endogenous CDK6, cells were fixed with methanol for 30 min at -20 °C. The fixed cells were further permeabilized with 0.1% Triton X-100 for 10 min at room temperature. After blocking in 3% BSA for 1 hr at room temperature or at 4 °C overnight, cells were incubated with appropriate primary antibodies at 4 °C overnight and secondary antibodies for 2 hrs at room temperature. Fluorescence images were acquired by Nikon ECLIPSE Ti microscopic system. The fluorescent intensity and the length of cilia or glutamylated axoneme were measured by NIS-Elements software (Nikon). The primary antibodies used for immunofluorescence are as follows: ARL13B (17711-1-AP; Proteintech, dilution 1:2000), acetylated tubulin (T7451, Sigma-Aldrich, dilution 1:5000), GT335 (AG-20B-0020-C100, AdipoGen Life Science, 1:4000), PolyE (AG-25B-0030-C050, AdipoGen Life Science, 1:1000), GLI-3 (AF3690, R&D Systems, dilution 1:500), RAB11FIP5 (14594-1-AP, Proteintech, dilution 1:1000), RAB11A (71-5300; Invitrogen, dilution 1:500), CDK7 (2916, Cell Signaling Technology, dilution 1:1000). E-Cadherin (24E10, Cell Signaling Technology, dilution 1:1000), Polycystin 2 antibody was provided by the Baltimore Polycystic Kidney Disease (PKD) Research and Clinical Core Center (dilution 1:1000).

For live-cell imaging experiments, FIP5-mcherry expressing RCTE stable cells were cultured in delta TPG dish (12-071-33, ThermoFisher Scientific Inc.) with 0.5 ml 10% FBS containing medium. At 24 hrs after plating cells at 20–30% confluence, RCTE cells were cultured in live-cell imaging culture chambers (Tokai Hit microscope stage top incubator) that set on Nikon ECLIPSE Ti microscope. For drug treatment, same volume (0.5 ml) medium with 2× concentration of abemaciclib (1 mM) was added to the petri dish. Images were taken every 10 min.

### Immunoprecipitation assay

Indicated plasmids were transfected into 293T cells. 48 hrs later, cell pellets were lysed in ice-cold lysis buffer (25mM Tris-HCl, pH 7.4, 150mM NaCl, 0.4% digitonin, 1 mM EDTA and protease inhibitors) for 30 min. The supernatant was collected by centrifugation for 20 min at 12,000g at 4 °C and further pre-cleared using protein-G Sepharose for 4 hrs. After removal of protein-G beads, the pre-cleared supernatant was incubated with protein-G beads and 2 μg of the indicated primary antibodies or IgG control overnight at 4 °C. After washing, the Sepharose beads were boiled in 1× SDS-PAGE loading buffer. The following commercially available antibodies were used for immunoprecipitation: Flag (8146, Cell Signaling Technology), mCherry (16D7, Invitrogen).

### RNA extraction and qPCR

For cell samples, total RNA was extracted from cells with the TRIzol reagent (Invitrogene). mRNAs were reverse transcribed using a high-capacity cDNA reverse transcription kit (Applied Biosystems, 4368814). Real-time PCR was performed using BlazeTaq SYBR Green qPCR Mix 2.0 (GeneCopoeia, QP031) with CFX384 Real-Time system (BIO-RAD).

The following primers were used: human *GAPDH* Fw 5′-TCCTGCACCACCAACTGCTT-3′ and Rv 5′-GTCTTCTGGGTGGCAGTGAT-3′; mouse *Gapdh* Fw 5′-TGCCCCCATGTTTGTGATG-3′ and Rv 5′-TGTGGTCATGAGCCCTTCC-3′; *Gli1* Fw 5′-GGTGCTGCCTATAGCCAGTGTCCTC-3′ and Rv 5′-GTGCCAATCCGGTGGAGTCAGACCC-3′; *Ptch1* Fw 5′-CTCTGGAGCAGATTTCCAAGG-3′ and Rv 5′-TGCCGCAGTTCTTTTGAATG-3′; *CDK5* Fw 5′-GATGATGAGGGTGTGCCGAG-3′ and Rv 5′-ATGAAGCCTGACGATGTTCT-3′; *Armc9* Fw 5′-TGAGCCAGACTACGGAGTTT-3′ and Rv 5′-GAGTCCAAGAATCCTGGAAG-3′.

### Immunohistochemistry

The sections of normal and ADPKD patients’ kidney tissues were gift from Dr. Peter Harris. WT and *Pkd1^RC/RC^* mice kidney tissues were fixed in formalin overnight at room temperature, paraffin embedded and sectioned. Paraffin blocks were cut into 5 mM sections Immunohistochemistry. The sections were dewaxed, rehydrated, pretreated, and treated with H_2_O_2_. The sections were incubated with CDK6 antibody (PA5-79027, Invitrogen, dilution 1:500) at 4°C overnight and secondary antibody (horseradish peroxidase-conjugated goat anti-mouse IgG) for 30 min at room temperature. After washing, the bound antibody was visualized using diaminobenzidine/H_2_O_2_ substrate, and the nuclei were restained with hematoxylin. The photographs were taken using Nikon Eclipse Ti2.

### Culture of primary renal tubular epithelial cells

WT and *Pkd1^RC/RC^* (*PKD1* hypomorphic allele, p.Arg3277Cys) C57BL/6J mice were anesthetized and kidneys were immediately removed and placed in cold (0°C) Hanks Balanced Salt Solution (HBSS). The renal capsule was removed, and the medulla was dissected and discarded. The remaining cortical tissue was minced and transferred to 10 mL HBSS containing Collagenase A (Roche, 50-100-3278) at 200 units/ml. Tubules were incubated at 37°C while rotating at 70 RPM for 35 minutes. Following digestion, density sedimentation with horse serum (Invitrogen, 31874) was used to inactivate enzymes and enrich for the proximal tubules. After 5 minutes sedimentation, the supernatant containing the proximal tubules was removed, transferred to another tube, and centrifuged for 7 minutes at 200 g. Tubules were washed once with 10 mL of HBSS and centrifuged at 200 g for 7 minutes. Tubules were then resuspended in Renal Epithelial Cell Basal Medium (ATCC, PCS-400-030), supplemented with Renal Epithelial Cell Growth Kit (ATCC, PCS-400-040) and 1% antibiotic/ antimycotic solution (Gibco, 15240062) and incubated at 37°C with 5% CO_2_.

### Metanephros Culture

Metanephroi were dissected from embryonic *Pkd1^RC/RC^*mice at E13.5 and placed on transparent Falcon 0.4-μm cell culture inserts in 12-well culture plate. 500 μl/well DMEM/F12-defined culture medium (supplemented with 2 mM L-glutamine, 10 mM HEPES, 5 μg/ml insulin, 5 μg/ml transferrin, 2.8 nM selenium, 25 ng/ml prostaglandin E, 3 ng/ml T3, 1% penicillin/streptomycin) was added under the culture inserts, and organ cultures were maintained in a 37°C humidified CO2 incubator for up to 4 days. To induce cystogenesis and test the effect of Abemaciclib, 10 μM Froskolin and Abemaciclib was added to the medium as indicated. Upon culturing and approximately 24 h later (1 d) and each day after (2 to 4 d), kidneys were photographed, and the images were acquired Nikon Eclipse Ti2.

Quantification of dilated tubule area was performed on captured images using NIS-Elements software. The wand tool was used to select a pixel within the image of a dilated tubule, which highlighted all of the pixels of similar density within the dilated tubule. This process was repeated within the dilation until the highlighted area comprised the entire dilation. For total kidney area, the freehand polygon tool was used to trace around the kidney (excluding the ureter), and the area within the tracing was determined. Fractional cyst area was calculated as total tubule dilation area divided by total kidney area.

### EdU cell proliferation Assay

For cell proliferation assay, cells were plated in 12-well plate with cover slide glass at desired density. Allow cells recover overnight to reach 60∼70% confluence. The cells then were incubated with 10 μM EdU for 3 hrs at 37 °C and followed by fixation and permeabilization steps. For fluorescent EdU detection, add reaction cocktail (20 mg/ml sodium ascorbate, 1 μM Alexa 594-azide, 4 mM CuSO4 in TBS) to each well and incubate for 30 minutes at room temperature and protect form light. After washing with 3% BSA (bovine serum albumin) in PBS, all slides were mounted with fluorescent mounting medium with DAPI to label nucleus and coverslipped. The EdU positive nucleus were quantified using Nikon ECLIPSE Ti microscopic system.

### Statistical information

For biological experiment analyses, data were analyzed using Prism 7 software (GraphPad) by two-tailed unpaired Student’s t-test, one-way or two-way ANOVA with Tukey’s post-hoc test. *P* < 0.05 was considered as significant (*\*P* < 0.05; *\*\*P* < 0.01; *\*\*\*P* < 0.001; n.s: not significant). Data are presented as mean ± s.d. or mean ± s.e.m. and sample size were indicated in each figure legend. No statistical methods were applied to pre-evaluate sample size. Ciliary length and fluorescent intensity measurements were performed at least twice with similar results, and a representative result is shown. qPCR of gene expression was performed with three or four biological replicates. No data were excluded from analysis. Samples in this study were not randomized. Blinding was not used for this study because the cell culture, sample preparation, reagents and experimental settings were kept consistent for each experiment.

## Acknowledgments

We thank Dr. Carsten Janke (Institute Curie, France) for sharing TTLL and CCP5 expressing constructs and Dr. Takanari Inoue (Johns Hopkins University) for sharing the cilia trapping system. This work was supported by the funding from National Institutes of Health (NIH) research grants (R01DK090038, R01DK099160, R01AG076469, and P30 center grant P30DK90728 to J. H.), Mayo Clinic Robert M. and Billie Kelley Pirnie Translational Polycystic Kidney Disease Center and Mayo Clinic Foundation (to J. H. and K.L.). Department of Defense grant (W81XWH2010214 to K.L.). This research was funded by a grant from the Polycystic Kidney Disease Foundation, pkdcure.org (to K.H). The Foundation had no role in study design, data collection and interpretation, or the decision to submit the work for publication.

## Data Availability

The authors declare that the data supporting the findings of this study are available within the manuscript. Any further relevant data are available from corresponding authors upon reasonable request.

## Expended View Figure Legends

**Figure EV1.**
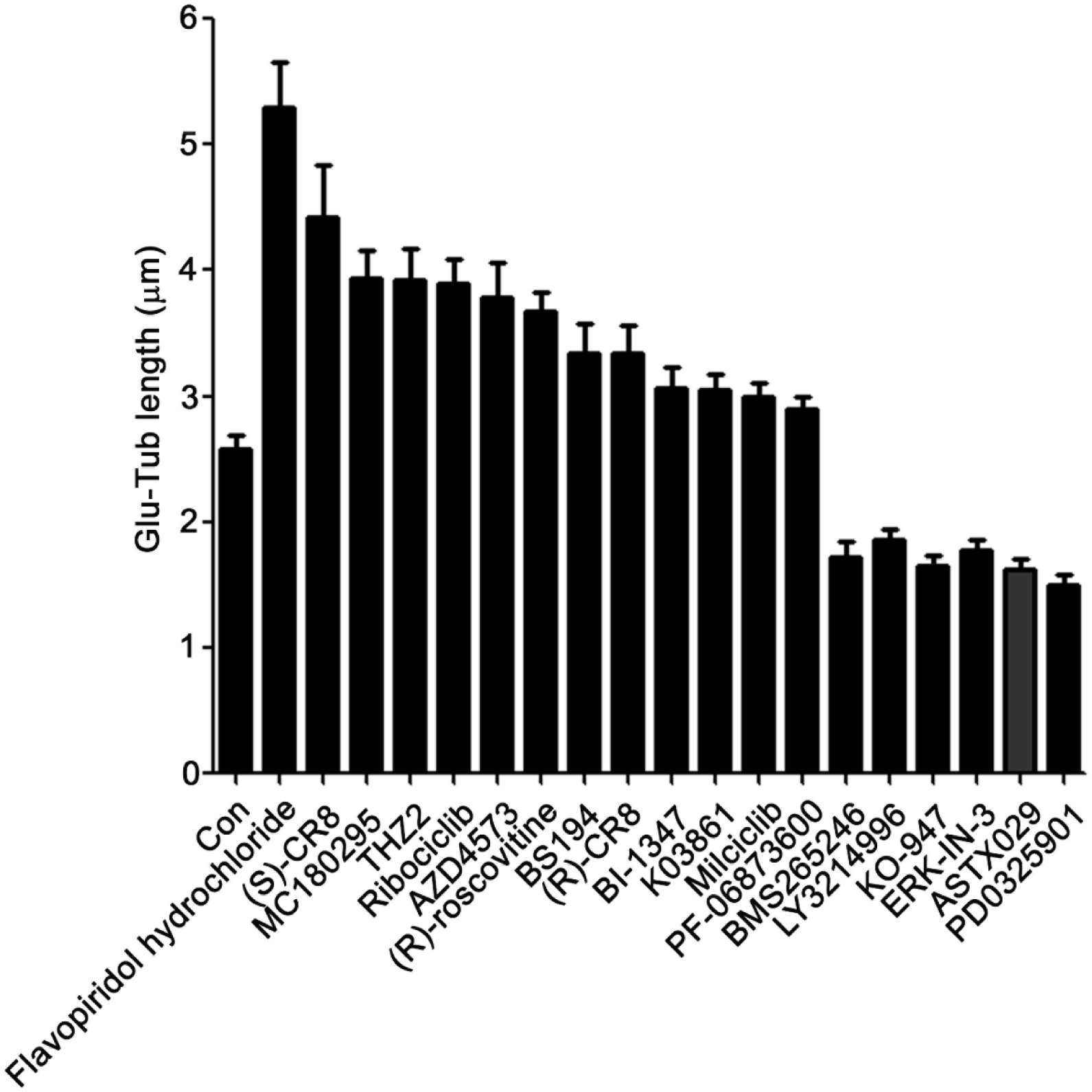
Kinase inhibitors screen to identify regulators of axoneme polyglutamylation. RPE-1 cell were treated with each kinase inhibitors for 24 hrs (1 μM) in serum free medium and the length of glutamylated axoneme were measured. The positive hits that affect axoneme polyglutamylation are presented. Data information: Quantified data are presented as mean ± s.e.m. N≥30 Cilia.

**Figure EV2.**
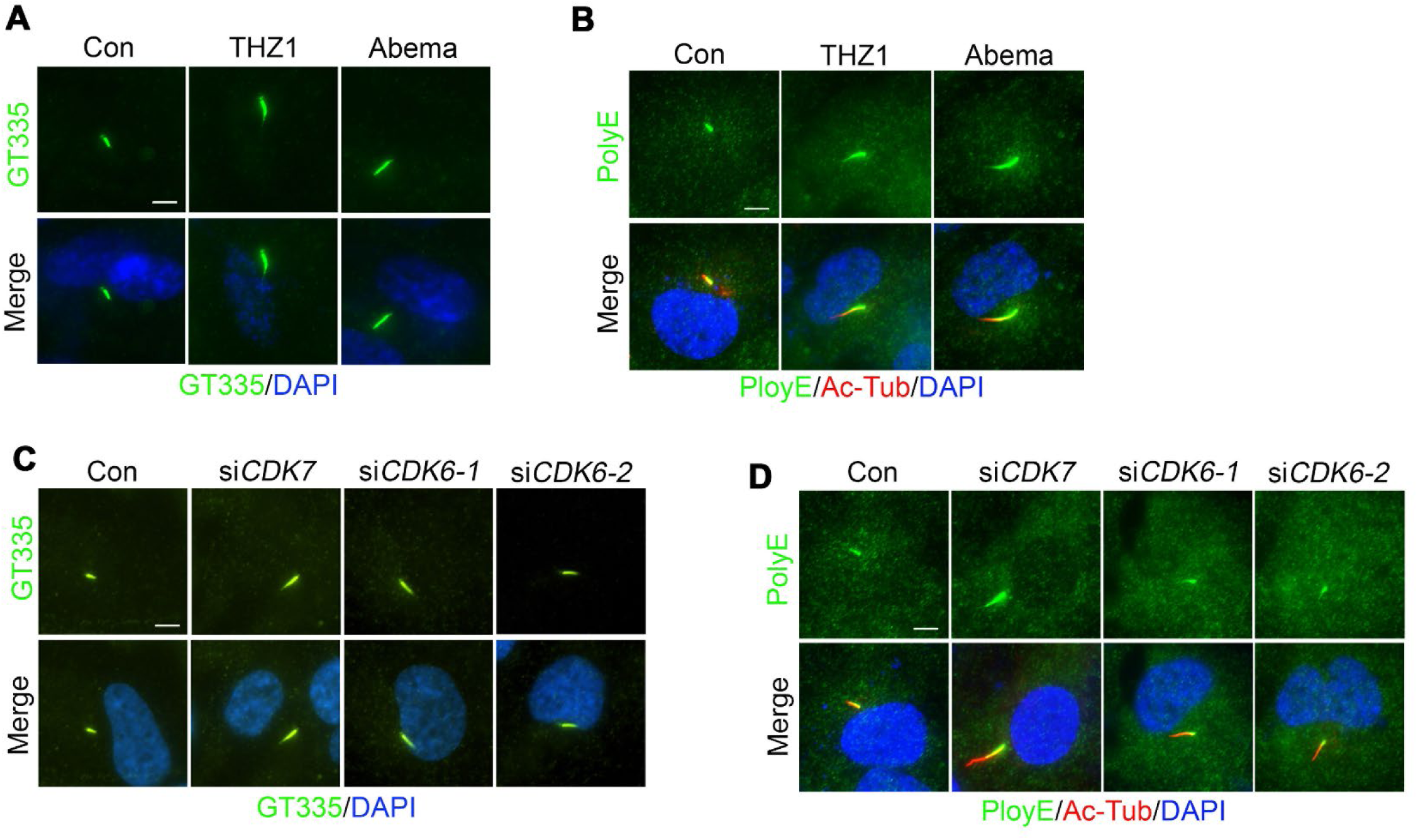
Inhibition of CDK7 or CDK6 does not affect tubulin glutamylation in cytoplasm. **A, B** THZ1 or Abemaciclib does not affect cytoplasmic tubulin glutamylation. **C, D** Knockdown of CDK7 or CDK6 does not affect cytoplasmic tubulin glutamylation. Data information: Scale bars: 5 μm.

**Figure EV3.**
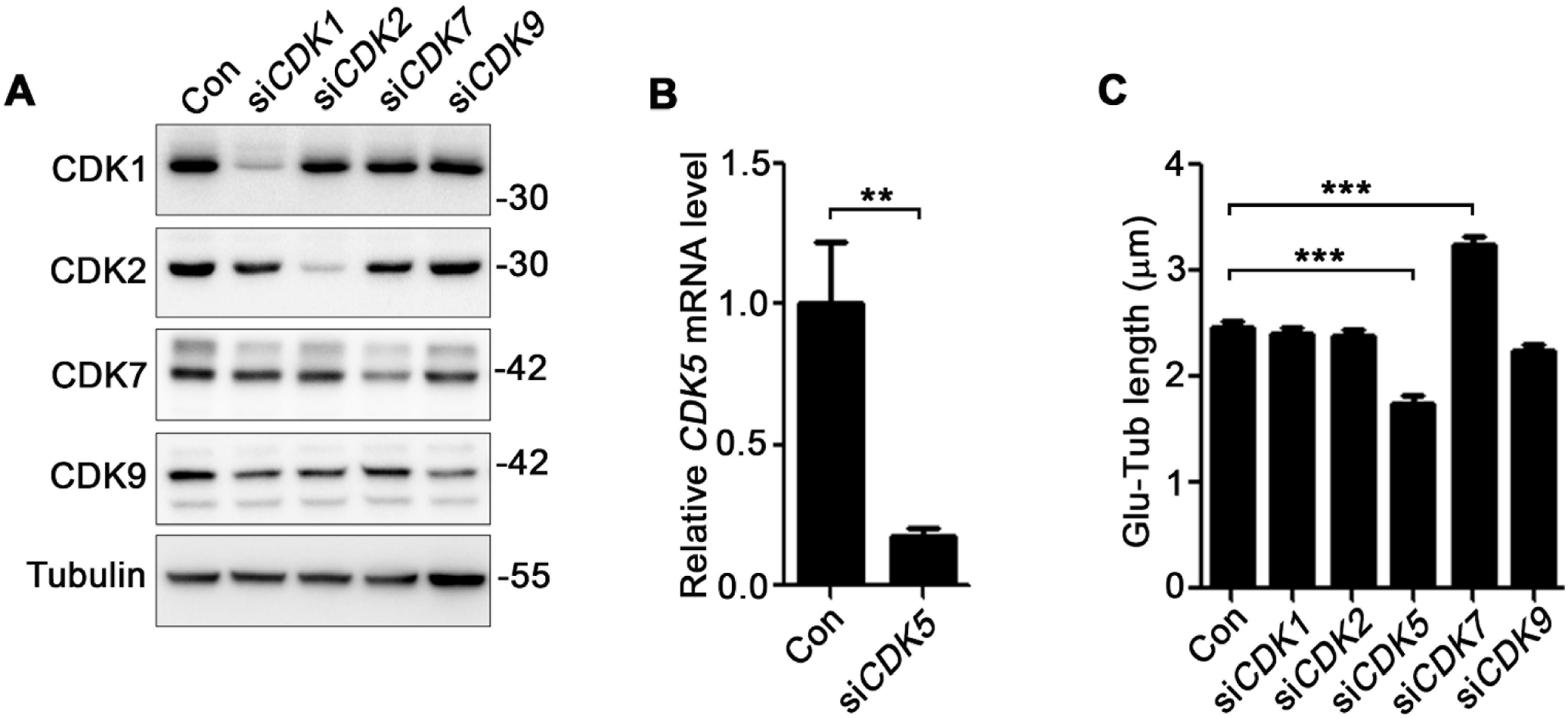
siRNA knockdown screen of CDK7 downstream CDKs. **A, B** The knockdown efficiency of indicated siRNAs in RPE-1 cells were accessed by western blotting (A) or qPCR (B). Quantified data are presented as mean ± s.d. N=3 independent experiments. Statistical analyses were performed by two-tailed unpaired Student’s t-test. **C** The effect of indicated CDKs knockdown on axoneme polyglutamylation in RPE-1 cells. Data information: Quantified data are presented as mean ± s.e.m. Statistical analyses were performed by one-way ANOVA analyses with Tukey’s post-hoc test for multiple comparisons. N≥50 Cilia. *\*\*P* < 0.01; *\*\*\*P* < 0.001.

**Figure EV4.**
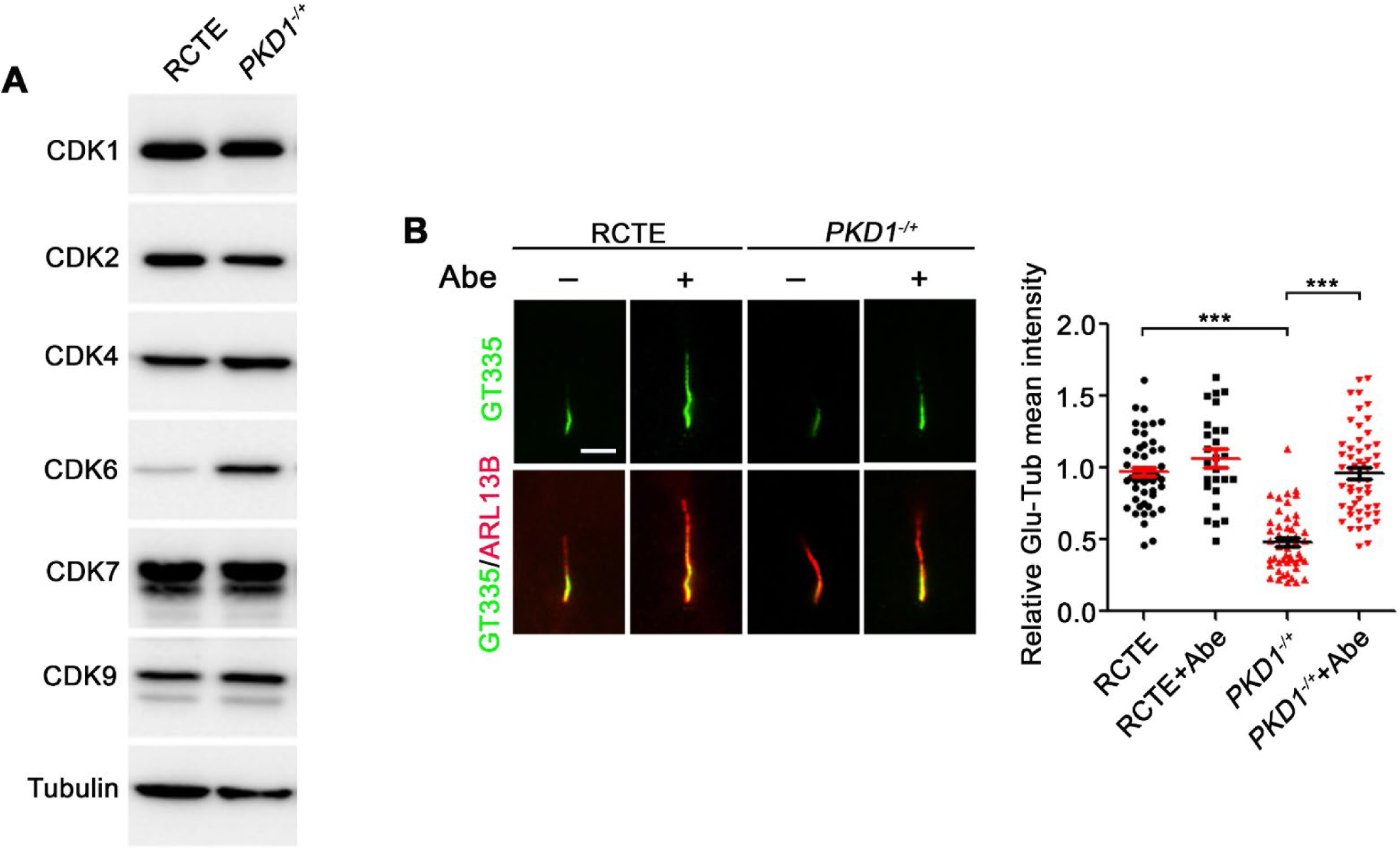
CDK6 upregulation and defective axoneme polyglutamylation in ADPKD patient-derived kidney tubular epithelial cells. **A.** The protein levels of indicated CDKs in RCTE cells and ADPKD patient-derived *PKD1^-/+^* kidney tubular epithelial cells were accessed by western blotting. **B.** Abemacilib (500 nM, 24 hrs) rescues axoneme polyglutamylation in ADPKD patient-derived kidney tubular epithelial cells. Data information: Scale bar: 2 μm. Quantified data are presented as mean ± s.e.m. Statistical analyses were performed by one-way ANOVA analyses with Tukey’s post-hoc test for multiple comparisons. N≥ 25 Cilia. \*\*\**P* < 0.001.

## Movie EV1. FIP5-positive vesicles show immediate enrichment upon Abemaciclib treatment

Live-cell time-lapse imaging of FIP5-mCherry expressing RCTE cells were performed during 2 hrs before and after abemaciclib treatment. Cells were cultured in 10% FBS containing medium. The time point of abemaciclib treatment was indicated by annotation (+Abemaciclib).

## Appendix Figure Legends

**Appendix Figure S1.**
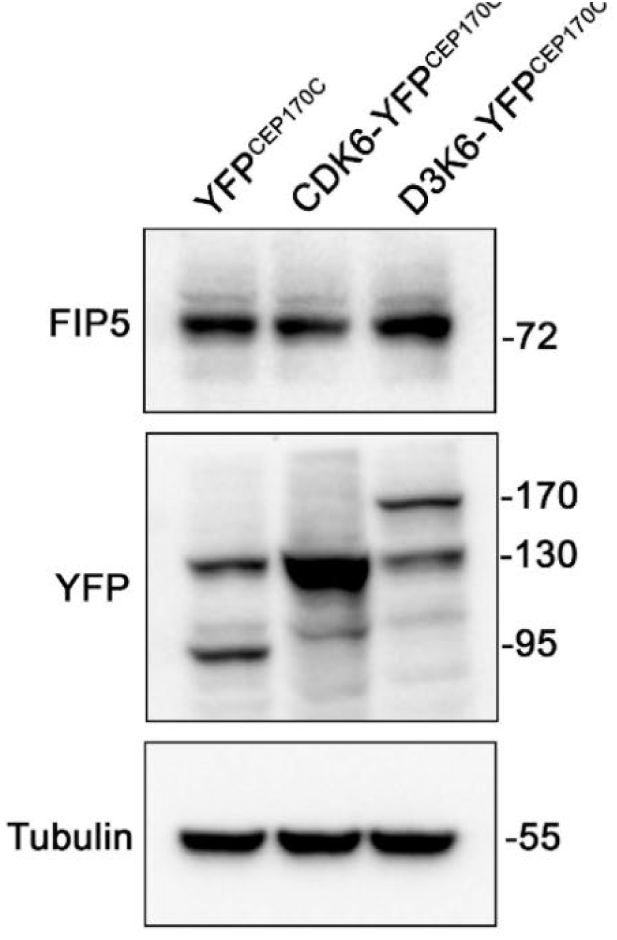
Ciliary CDK6 does not affect the protein level of FIP5. YFP control, CDK6 or CyclinD3-CDK6 were overexpressed at cilia base. The protein level of FIP5 was accessed by western blotting.

**Appendix Figure S2.**
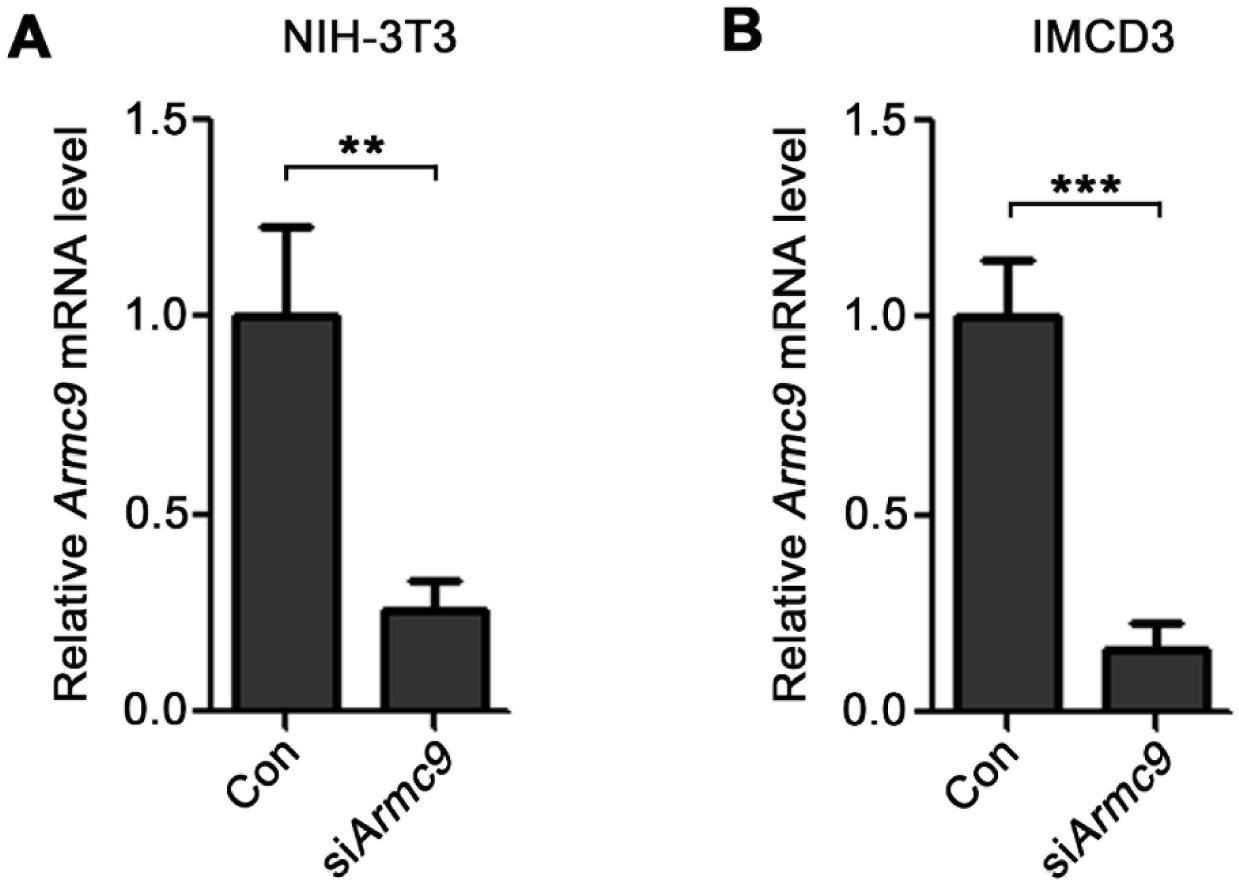
The knockdown efficiency of *Armc9* siRNA. **A, B** The knockdown efficiency of *Armc9* siRNA in NIH-3T3 (A) and IMCD3 (B) cells were accessed by qPCR. Data information: Quantified data are presented as mean ± s.d. Statistical analyses were performed by unpaired student t-test. N=3 independent experiments. *\*\*P* < 0.01; *\*\*\*P* < 0.001.

**Appendix Figure S3.**
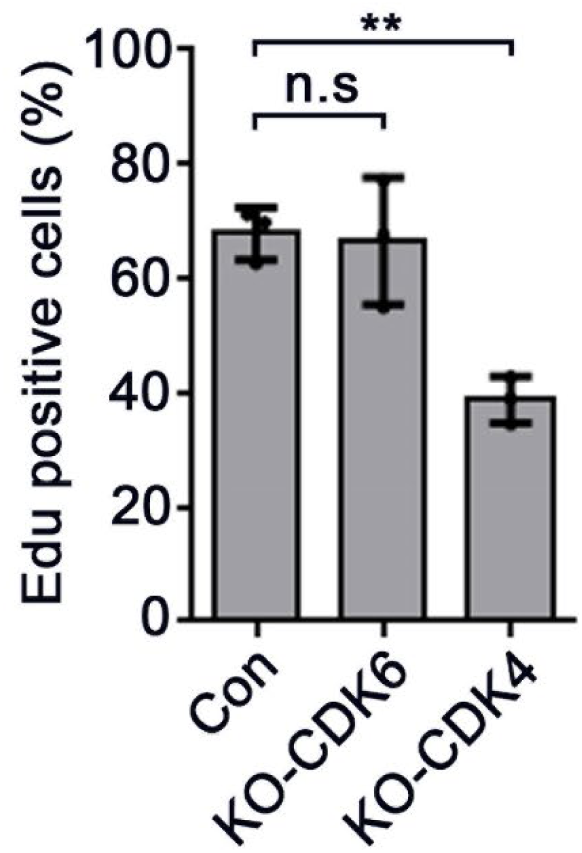
CDK6, but not CDK4, is dispensable for cell proliferation in RCTE cells. The cell proliferation of *WT*, *CDK6^-/-^* and *CDK4^-/-^* RCTE cells were accessed by EdU incorporation assay. Data information: Quantified data are presented as mean ± s.d. Statistical analyses were performed by one-way ANOVA analyses with Tukey’s post-hoc test for multiple comparisons. N=3 independent experiments. n.s: not significant; *\*\*P* < 0.01.

**Appendix Table S1.**
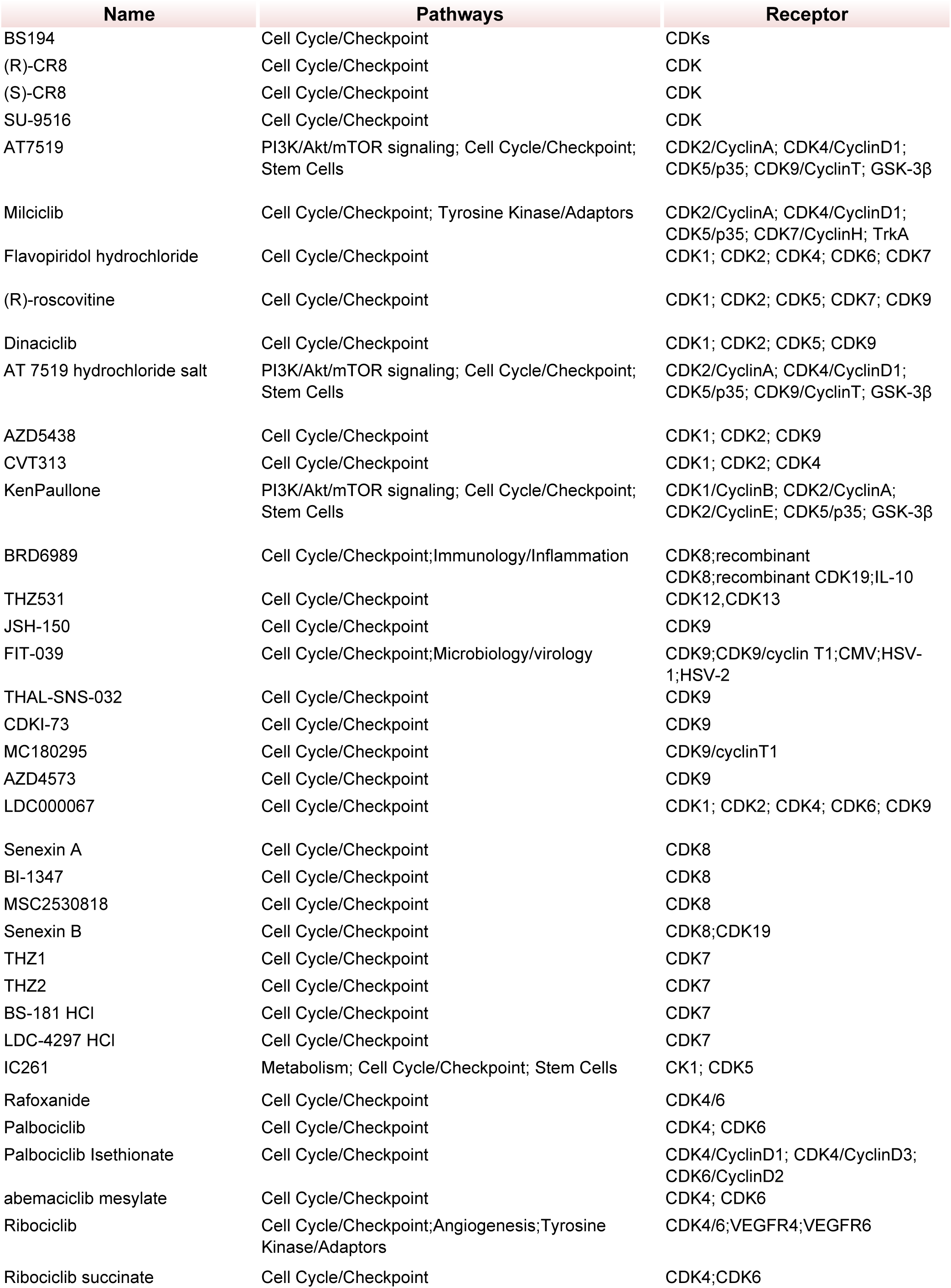
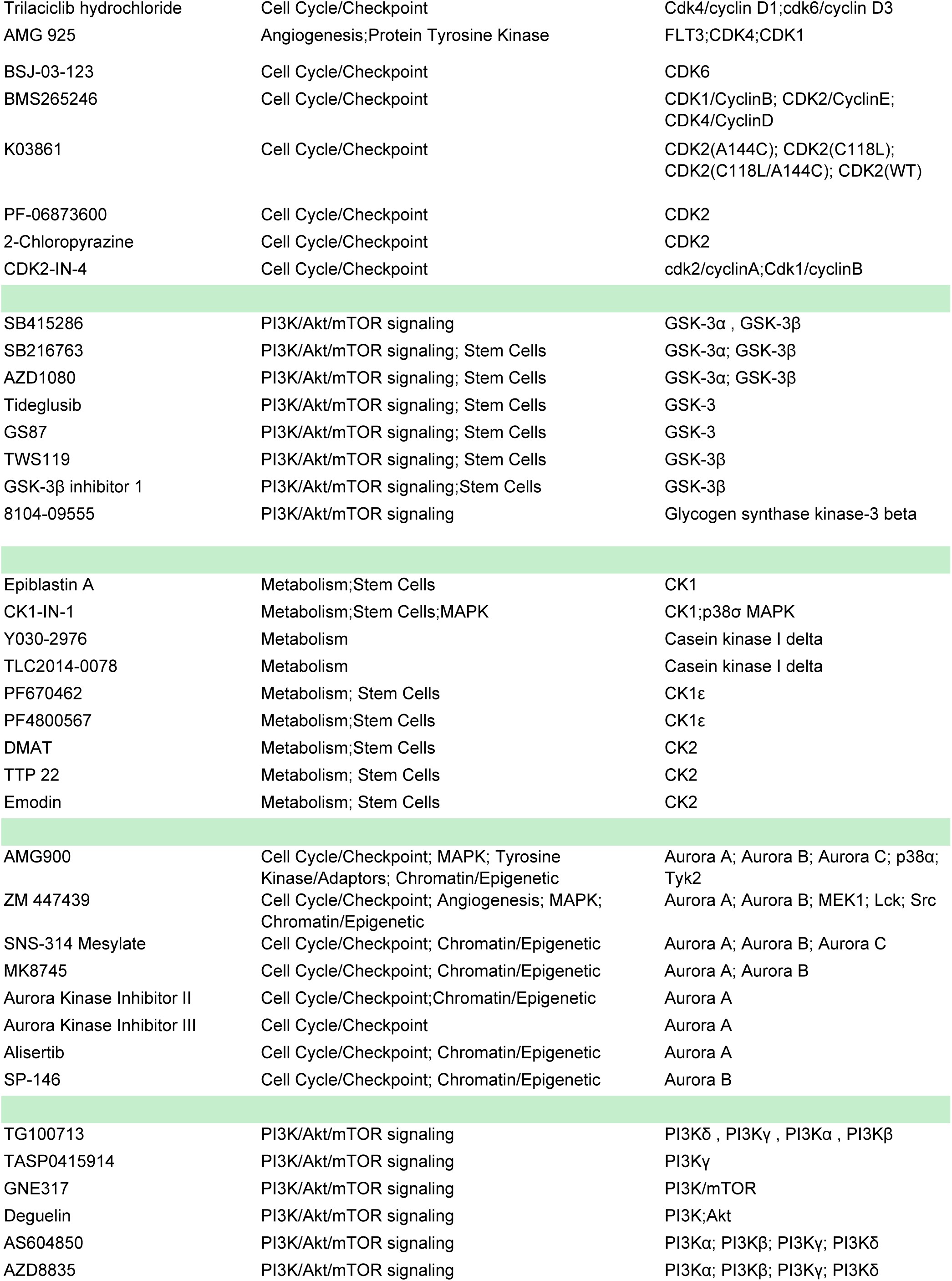
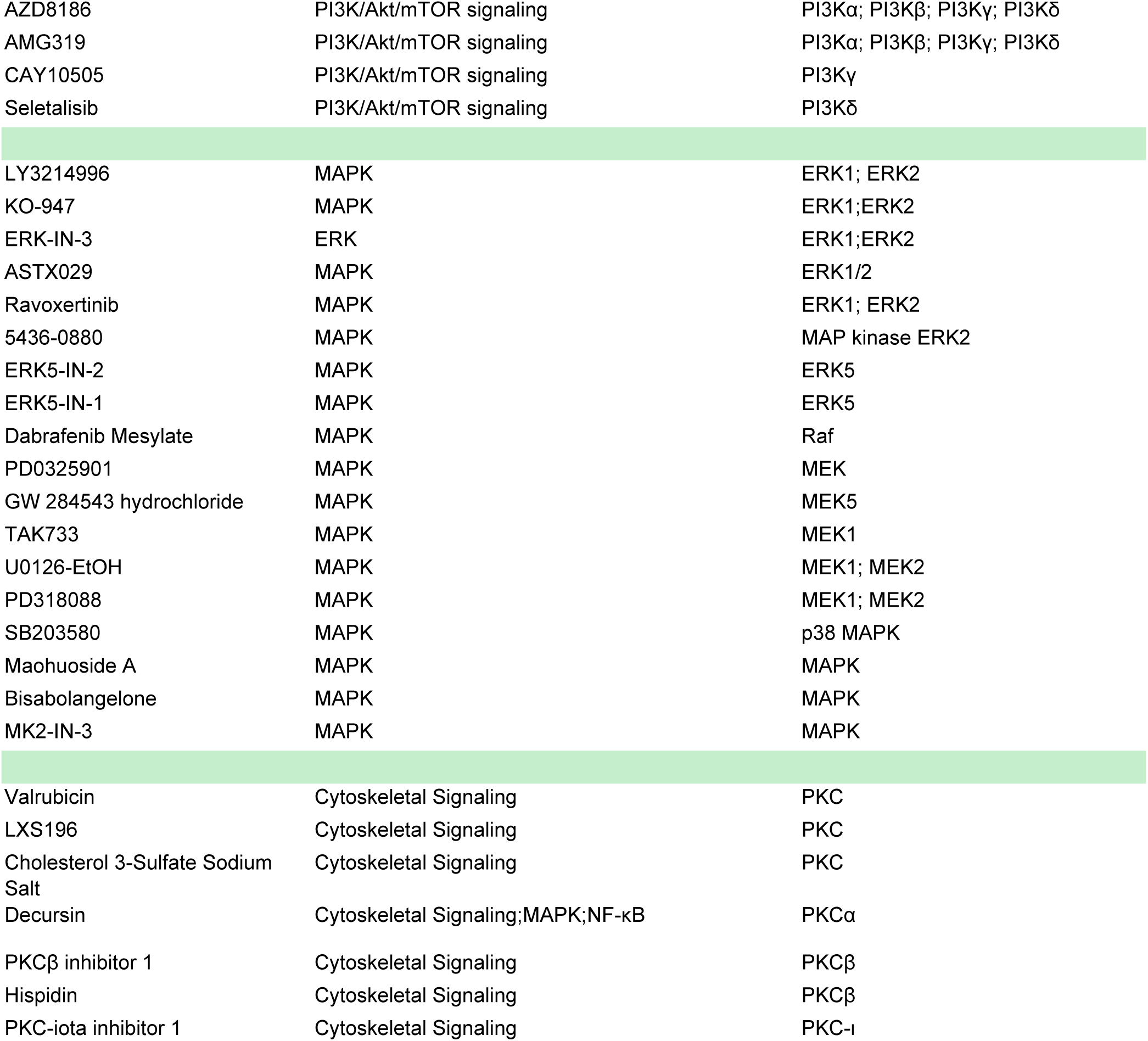
List of kinase inhibitors screen. The name, targeted pathway(s), and targeted receptor(s) of tested kinase inhibitors are listed in this table.

